# Assessing simulation-based supervised machine learning for demographic parameter inference from genomic data

**DOI:** 10.1101/2025.04.07.647546

**Authors:** Arnaud Quelin, Frédéric Austerlitz, Flora Jay

**Author notes:** Corresponding author: Arnaud Quelin. These authors contributed equally to this work (Frédéric Austerlitz, Flora Jay).

## Abstract

The ever-increasing availability of high-throughput DNA sequences and the development of numerous computational methods have led to considerable advances in our understanding of the evolutionary and demographic history of populations. Several demographic inference methods have been developed to take advantage of these massive genomic data. Simulation-based approaches, such as approximate Bayesian computation (ABC), have proved particularly efficient for complex demographic models. However, taking full advantage of the comprehensive information contained in massive genomic data remains a challenge for demographic inference methods, which generally rely on partial information from these data. Using advanced computational methods, such as machine learning, is valuable for efficiently integrating more comprehensive information. Here, we showed how simulation-based supervised machine learning methods applied to an extensive range of summary statistics are effective in inferring demographic parameters for connected populations. We compared three machine learning (ML) methods: a neural network, the multilayer perceptron (MLP), and two ensemble methods, random forest (RF) and the gradient boosting system XGBoost (XGB), to infer demographic parameters from genomic data under a standard isolation with migration model and a secondary contact model with varying population sizes. We showed that MLP outperformed the other two methods and that, on the basis of permutation feature importance, its predictions involved a larger combination of summary statistics. Moreover, they outperformed all three tested ABC algorithms. Finally, we demonstrated how a method called SHAP, from the field of explainable artificial intelligence, can be used to shed light on the contribution of summary statistics within the ML models.

## Introduction

Deciphering the demographic history of natural populations through the analysis of genetic polymorphism data represents an ongoing challenge in the field of population genetics due to the complexity of these data and their underlying processes. A better understanding of past demographic events can provide, for example, insights on specific questions such as the impact of anthropogenic processes (Pujolar et al., 2017; Dong et al., 2021) or climatic events (Bai et al., 2018; Fedorov et al., 2020) on population dynamics. It can also be helpful in conservation biology (Der Sarkissian et al., 2015; Abascal et al., 2016).

The exploration of genetic diversity also sheds light on the intricate dynamics of human populations (Cavalli-Sforza et al., 1994; Henn et al., 2012; Schraiber & Akey, 2015). Analysing genomic polymorphisms in the genomes of modern human populations enables us to infer past demographic and evolutionary events on different timescales, including colonization, separation, migration and expansion events, as well as adaptive processes. As such, reconstructing the demographic history of populations is essential to disentangle the effects of demography from those of selection (Akey et al., 2004; Johri et al., 2023; Lohmueller, 2014).

Taking full advantage of the information contained in a large number of genomic data remains a methodological and computational challenge. Several kinds of inference methods have been developed to infer past demographic events from these genomic data (Beichman et al., 2018; Marchi et al., 2021). Some of these methods are based on the sequential Markovian coalescent (SMC) model (McVean & Cardin, 2005; Marjoram & Wall, 2006; Li & Durbin, 2011; Sheehan et al., 2013; Schiffels & Durbin, 2014; Terhorst et al., 2017). They use the pattern of diversity along the genomes to estimate the inverse coalescent rate through time, which is a proxy for past population sizes under specific assumptions (Chikhi et al., 2018). Other methods use the length of regions that are identical-by-state (IBS) or identical-by-descent (IBD) between haplotypes (Palamara et al., 2012; Harris & Nielsen, 2013; Browning & Browning, 2015). Finally, some methods rely solely on the site frequency spectrum (Gutenkunst et al., 2009; Liu & Fu, 2015). Unlike the other methods, they do not take into account the linkage disequilibrium among SNPs.

Approximate Bayesian computation (ABC) is a simulation-based method used when the likelihood function is unavailable or too expensive to compute. It initially emerged to tackle population genetics issues (Tavaré et al., 1997; Pritchard et al., 1999; Beaumont et al., 2002) and was then used in various domains such as ecology (Jabot & Lohier, 2016; Wood, 2018), epidemiology (Tanaka et al., 2006; McKinley et al., 2018), system biology (Toni et al., 2008; Liepe et al., 2014) or linguistics (Thouzeau et al., 2017, 2022). This framework appears particularly efficient when dealing with complex demographic models (Beaumont, 2010; Bertorelle et al., 2010). It relies on the comparison of summary statistics computed on the genomic data set under investigation and on genomic data sets simulated under a given demographic model, for which the parameters are generally drawn in uninformative prior distributions. In the classical rejection sampling approach, the posterior distributions of the parameters are estimated by keeping only the simulations yielding summary statistics close to those from the real data (Tavaré et al., 1997). More elaborate methods were subsequently designed (Blum & Francois, 2010; Estoup et al., 2012).

More recently, simulation-based approaches combined with supervised machine learning algorithms were developed for demographic inference (Schrider & Kern, 2018; Collin et al., 2021; Korfmann et al., 2023). Some of these approaches are based on random forests (Pudlo et al., 2016; Raynal et al., 2019; Ghirotto et al., 2021), while others use neural networks (Sheehan & Song, 2016; Mondal et al., 2019; Sanchez et al., 2021). As machine learning (ML) methods in demographic inference have gained ground in recent years, the focus has not been on comparing their characteristics and performance.

In this study, we focused on the inference of demographic parameters using coalescent simulations for models in which an ancestral population splits into two at some point in the past, with these populations remaining connected by migration. We implemented three machine-learning methods aiming at inferring demographic parameters from a large number of summary statistics computed on genomic data. These methods were based, respectively, on random forest (Breiman, 2001), XGBoost (Chen & Guestrin, 2016), and multilayer perceptron (Haykin, 1994). To our knowledge, it was the first time that XGB was used in this context.

After comparing their overall performance and dissecting the impact of demographic parameters on the results, we analyzed which summary statistics contributed most and introduced a method to evaluate their influence on predictions.

## Methods

Broadly, our approach consists of (i) producing many simulated datasets under a demographic model, (ii) computing summary statistics on these simulated datasets, and (iii) using these summary statistics as a training dataset for three machine-learning methods.

### Demographic models

We first considered a standard model of isolation with migration (IM) (Fig. 1A), where an ancestral population splits into two populations at a given time (*Split_time*). We assumed that, right after this splitting event, a continuous and symmetrical migration rate (*Migration_rate*), defined as the proportion of individuals moving from one population to the other per generation, was maintained between the two populations until the present. These two populations were assumed to have distinct effective population sizes (*N_current_1* and *N_current_2*), themselves distinct from the ancestral effective population size (*N_ancestral*). In this model, effective population sizes were assumed to be constant except for the change at split time.

**Fig. 1.**
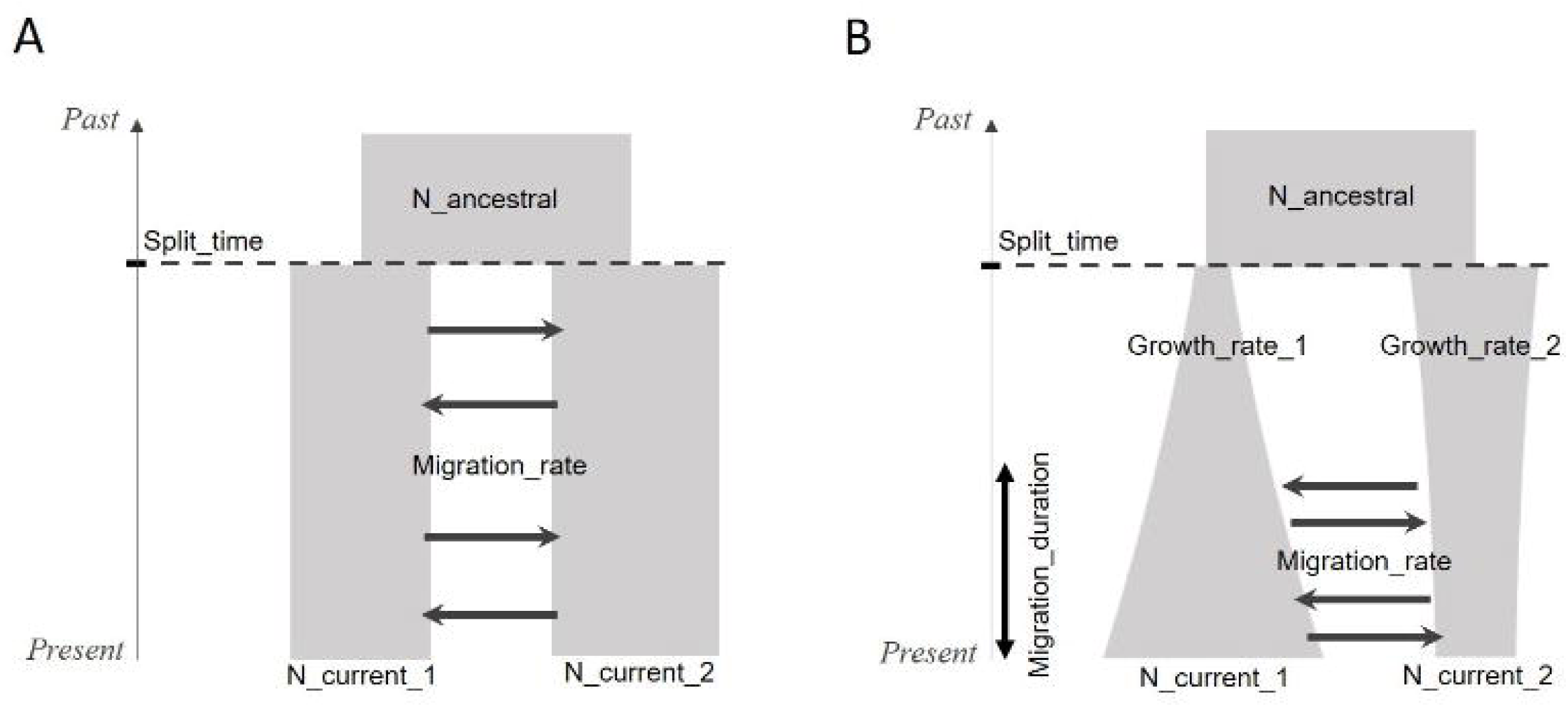
Demographic models with the varying demographic parameters. A-Isolation with migration (IM) model. B – Secondary contact with varying population sizes (SC) model. Both populations can either grow or decline.

We then considered a more complex model of secondary contact with varying population sizes (SC) (Fig. 1B). In this model, the effective population sizes of the two populations resulting from the split were no longer assumed to be constant but instead to increase or decrease exponentially, with different growth rates for the two populations. Furthermore, the populations did not undergo gene flow immediately after their separation: migration started at a later stage and lasted until the present time. This allowed us to assess the performances of the ML models on a more complex model.

### Simulations

We generated 10,000 simulated data sets using *msprime* (Kelleher et al., 2016; Baumdicker et al., 2021) for each of the two demographic models. For the simulated data sets under the IM model, the five parameters were drawn in uniform prior distributions: Uniform[1;5,000] for *Split_time*, corresponding to the number of generations since the split occurred; Uniform[100;10,000] for *N_ancestral*, *N_current_1*, and *N_current_2*; and Uniform[0;0.001] for *Migration_rate*. While other prior distributions could be considered, we opted for uniform distributions to ensure an even exploration of the parameter space. We note that by setting the lower bound of the *Split_time* prior to 1 generation, predictions can be particularly difficult in such extreme scenarios. For each parameter set, we simulated 20 independent neutral loci for a sample of 10 diploid individuals in each population. Each locus was a genomic sequence of length 2Mb, with a constant recombination rate of 1.0×10^-8^ per bp per generation and a constant mutation rate of 1.25×10^-8^ per bp per generation, which are standard values for human genomes (Campbell et al., 2012; Kong et al., 2012; Scally & Durbin, 2012; Schiffels & Durbin, 2014).

For the SC model, the five parameters already present in the IM model were drawn from uniform prior distributions that differed to some extent from the priors of the IM model: Uniform[100;5,000] for *Split_time*; Uniform[1,000;5,000] for *N_ancestral*, *N_current_1*, and *N_current_2*; and Uniform[0;0.005] for *Migration_rate*. In addition to the IM model, each population experienced exponential growth with per generation rates *Growth_rate_1* and *Growth_rate_2*, drawn independently in a uniform distribution: Uniform[-0.001;0.002]. Finally, the migration duration parameter was defined as the ratio between the number of generations during which migration occurred and the total number of generations since the split time. It was drawn in a uniform distribution Uniform[0,1], where 0 corresponded to a case with no migration and 1 to a case of continuous migration since the split.

Among the 10,000 simulated data sets, 5,000 were used as a training data set (used to train the models), 2,500 as a validation data set (used to tune hyperparameters, i.e., the parameters of the models set before the training process), and 2,500 as a test data set (used to evaluate the final performance of the trained models).

### Summary statistics

The summary statistics that we computed on the genomic data belonged to 11 classes: eight classes were computed within populations and three among populations. Note however that the within-population statistics were computed for each population sample separately, and also on the pooled sample combining the two samples. The number of individual summary statistics varied considerably among the different summary statistics classes. Note that during training, these individual features were considered independently from the class to which they belong.

The within-population statistics classes were:

i. *S*: the proportion of segregating sites.
ii. *D*: Tajima’s *D* statistic (Tajima, 1989).
iii. *PI*: The mean and the standard deviation of the expected heterozygosity across segregating sites (Nei & Li, 1979).
iv. *WinH*: the mean and the standard deviation of the haplotypic heterozygosity computed for non-overlapping windows of size 50 kb.
v. Site frequency spectrum (*SFS*) statistics: The percentage of single-nucleotide polymorphisms (SNPs) where the derived allele is present at frequency *i*, for all *i* in [1, …, 2*n*], where *n* is the sample size, and the standard deviation of the distances between two adjacent SNPs with the derived allele at frequency *i* (Boitard et al., 2016; Jay et al., 2019).
vi. *LD*: mean and standard deviation of the squared correlation *r*^2^ of all pairs of SNPs in a given bin of distance. Following Boitard et al. (2016) and Jay et al. (2019), 19 bins of distance with means ranging from 282 bp to 1.4 Mb were considered.
vii. *IBS*: Deciles of segment length distribution completely identical between *m* haplotypes for *m* = 2, 4, 8, and 16 (Boitard et al., 2016; Jay et al., 2019; inspired by Harris & Nielsen, 2013).
viii. *AFIBS*: For each SNP, the *AFIBS* segment refers to the region surrounding the SNP that remains identical across all haplotypes containing the derived allele at that particular SNP (Theunert et al., 2012). We computed the mean and the standard deviation of the length of *AFIBS* segments for the SNPs of frequency *i* for all *i* in [1, …, 2*n*]. The among-population statistics classes were:
ix. *Fst*: the fixation index (Wright, 1950).
x. *Dxy*: genetic divergence between the two populations (Nei & Li, 1979).
xi. *JSFS*: Joint site frequency spectrum (Wakeley & Hey, 1997): The percentage of SNPs where the derived allele is present at a frequency *i* in population 1 and *j* in population 2.

The statistics of the classes *S*, *D*, *PI*, *Fst*, *Dxy,* and *JSFS* were calculated using the *tskit* package (Kelleher et al., 2016), while those for classes *WinH, SFS, LD, IBS* and *AFIBS* were computed using scripts adapted from Jay et al. (2019) in Python 3.9.7. All summary statistics were computed for each independent locus. We then computed the mean, median, and variance across loci of these values for each simulation, yielding a total of 3024 summary statistics (Table S1).

### Parameter inference methods

We explored the effectiveness of three regression methods to infer the demographic parameters: random forest (RF), gradient boosting system XGBoost (XGB), and multilayer perceptron (MLP). All models underwent optimization of their hyperparameters using a validation set. The CPU times used to train and predict these three methods are shown in Fig. S1.

RF (Breiman, 2001) is an ensemble method based on the construction of multiple decision trees. This method reduces the uncertainty of the prediction by averaging many predictions from the individual trees. The hyper-optimization procedure led us to set the number of trees between 200 and 300, with depths ranging from 15 to 20 (the exact values depended upon the demographic parameters).

While RF uses independent predictors, XGB (Chen & Guestrin, 2016) uses a sequential approach where each new tree is adjusted to correct errors in the existing model. This method assigns higher weights to observations that were poorly predicted in earlier iterations, thus prioritizing their correction in the construction of subsequent decision trees. For this second ensemble learning technique, the tuning of hyperparameters on the validation set led us to set the number of trees to 100, with a depth of seven.

Finally, MLP (Haykin, 1994) is a neural network method made up of fully connected layers. We trained a separate network for each demographic parameter, although a multi-output MLP could also be used. Depending on the demographic parameters, a hyper-optimization preliminary study on the validation set led us to set the number of neuron layers to six or seven (Table S2). For all models, summary statistics were standardized before training and the loss function was the mean square error (MSE). Unlike RF or XGB, MLP can lead to predictions outside the prior intervals. In that case, the prediction was set to the closest bound of the prior interval. All MLP models incorporate the ReLU function as an activation function (Agarap, 2019). We also performed a preliminary study on the impact on prediction quality of the introduction of an L1 regularization penalty, which led us not to use this option (Fig. S2).

We implemented the RF and MLP methods using the scikit-learn python library (Pedregosa et al., 2011), and the XGBoost library for the XGB method (Chen & Guestrin, 2016).

### Feature contributions

We used permutation feature importance (PFI) to gauge the contribution of each feature (i.e., each summary statistic) to the performance of a specific model. It consists in randomly shuffling the values of a given feature among replicates and observing the subsequent degradation of the model’s performance. By breaking the feature-target relationship, it allows to determine the extent to which the model depends on this particular feature. In this study, we used the Permutation importance function from the sickit-learn library.

Moreover, we used Shapley values to assess the extent to which each summary statistic impacts the model’s predictions for each demographic parameter, measuring both the direction and magnitude of their contributions. Shapley values first appeared in a context of cooperative game theory (Shapley, 1953) to estimate the contribution of individual players to the outcome of a game. In the ML context, the players are represented by the features and the cooperation is represented by combinations of features. We used the Python SHAP library (Lundberg & Lee, 2017), which provides visual representations of the influence of the feature value on the outcome.

### Comparison with ABC methods

The three regression methods were compared with several ABC algorithms from the ‘abc’ package (Csilléry et al., 2012), namely ABC rejection, ABC loclinear and ABC neuralnet, tested on the same simulation set. In addition to a rejection step, the last two methods apply corrections based on a local linear regression and a neural network, respectively. Their efficiency was computed on the same 2,500 samples from the test data set used to assess the regression methods. For each of these 2,500 samples, we kept the simulations closest to the target among the 7,500 simulations not used in the test data set, for tolerance thresholds ranging from 5×10^-4^ to 5×10^-2^ (Table S3).

For the loclinear method, we selected the summary statistics with the highest correlation with the demographic parameters, in order to comply with the limitations of this method on the maximal number of summary statistics. Thus, we retained up to 300 statistics in the case of the 5×10^-2^ tolerance threshold. Similarly, to follow neuralnet specifications, we kept the 300 statistics with the highest correlation with the target parameter while considering a network with three neurons in its hidden layer. For all three algorithms, we then used either the mean or the median of the posterior as a point estimate.

## Results

### Accuracy of the machine-learning method for parameter estimation under the IM model

After training the three ML methods for each demographic parameter of the standard IM model on the training data set, we evaluated their performances on the test data set. We compared the predictions of these three ML methods using different metrics (Table 1). In the following, *N_current* refers to results for population 1. Since the model is symmetric and both population sizes have the same prior, the results obtained for *N_current_1* and *N_current_2* are similar (see Table S4 for population 2).

**Table 1.** Prediction errors on the test data set of the three ML methods, for each demographic parameter of the standard IM model. RMSE: Root Mean Square Error; MAE: Mean Absolute Error; NMAE: Mean Absolute Error normalized by the range of the parameter. The lowest errors for each parameter are indicated in bold.

| Demographic parameter | Method | RMSE | MAE (*se*) | NMAE (*se*) |
| --- | --- | --- | --- | --- |
| Split_time | RF | 7.28E+02 | 5.22E+02 (1.02E+01) | 1.04E-01 (2.03E-03) |
|  | XGB | 6.76E+02 | 4.67E+02 (1.02E+01) | 9.34E-02 (1.95E-03) |
|  | MLP | **6.35E+02** | **3.85E+02** (9.77) | **7.71E-02** (2.01E-03) |
| Migration_rate | RF | 1.32E-04 | 8.82E-05 (1.97E-06) | 8.82E-02 (1.97E-03) |
|  | XGB | 1.22E-04 | 8.00E-05 (1.84E-06) | 8.00E-02 (1.84E-03) |
|  | MLP | **1.21E-04** | **7.52E-05** (1.91E-06) | **7.52E-02** (1.91E-03) |
| N_ancestral | RF | 1.84E+02 | 1.29E+02 (2.61) | 1.29E-02 (2.61E-04) |
|  | XGB | **1.81E+02** | 1.30E+02 (2.51) | 1.30E-02 (2.52E-04) |
|  | MLP | 1.89E+02 | **1.27E+02** (2.81) | **1.27E-02** (2.81E-04) |
| N_current | RF | 6.55E+02 | 4.27E+02 (9.95) | 4.27E-02 (9.96E-04) |
|  | XGB | 6.09E+02 | 3.90E+02 (9.36) | 3.91E-02 (9.36E-04) |
|  | MLP | **5.98E+02** | **3.74E+02** (9.32) | **3.75E-02** (9.33E-04) |

Root mean square error (RMSE) and mean absolute error (MAE) are commonly used measures, and the choice between them depends on the specificities of the issue to be tackled (Chai & Draxler, 2014). The squaring operation in RMSE penalizes large errors more heavily, which is particularly appropriate when avoiding outlier prediction is critical. For three of the four demographic parameters, *Split_time*, *Migration_rate* and *N_current*, MLP performed better than RF and XGB for these two metrics. Conversely, for the *N_ancestral* parameter, the XGB model performed better according to the RMSE metric, while the MLP model performed better according to the MAE metric.

The MAE metric presents the advantage of directly reporting the average magnitude of the errors. *Split_time* was the demographic parameter for which the MLP model outperformed the two other models by the largest margin: the MAE of RF and XGB were 522 and 467 generations respectively, i.e., 36% and 21% higher than the MAE of MLP, which was only of 385 generations. For the migration rate, which ranged between zero and 10^-3^ per generation in the simulations, MLP achieved an average MAE of around 7.5×10^-5^ per generation, compared with 8.0×10^-5^ and 8.8×10^-5^, respectively, for the XGB and RF models, i.e., a higher MAE by 6% and 17%, respectively. For ancestral and current effective population sizes, the lowest average errors were also achieved by the MLP models. For ancestral effective population size, MLP outperformed only marginally the other methods, with a mean error for MLP of 127 individuals compared with 129 and 130 individuals, respectively, for the RF and XGB models. The difference was more substantial for the current effective population size: for this parameter, the MAE of RF and XGB were higher by 14% and 4%, respectively, than the MAE of 375 of the MLP.

The NMAE represents a normalized version of the mean absolute error (MAE), in which the error for a given parameter is divided by the difference between the minimum and maximum bounds of the prior of this parameter. This allowed us to have a comparison between the estimates of all parameters, whatever their range. *N_ancestral* had the lowest NMAE (0.0127). It was thus the parameter that could be estimated with the highest accuracy, followed by *N_current* (NMAE of 0.0375). *Migration_rate* and *Split_time* parameters, with respective NMAE of 0.0752 and 0.0771, showed the lowest accuracy.

Observing predicted versus observed values allowed us to obtain more detailed information about the predictions for each parameter (Fig. 2; Fig. S3). RF and XGB methods showed both a downward bias for high values of splitting time and migration rate, in contrast to MLP (Fig. 2). We also observed significant upward biased predictions for the RF model for low migration rates, leading to a large overall error for this method on this demographic parameter.

**Fig. 2.**
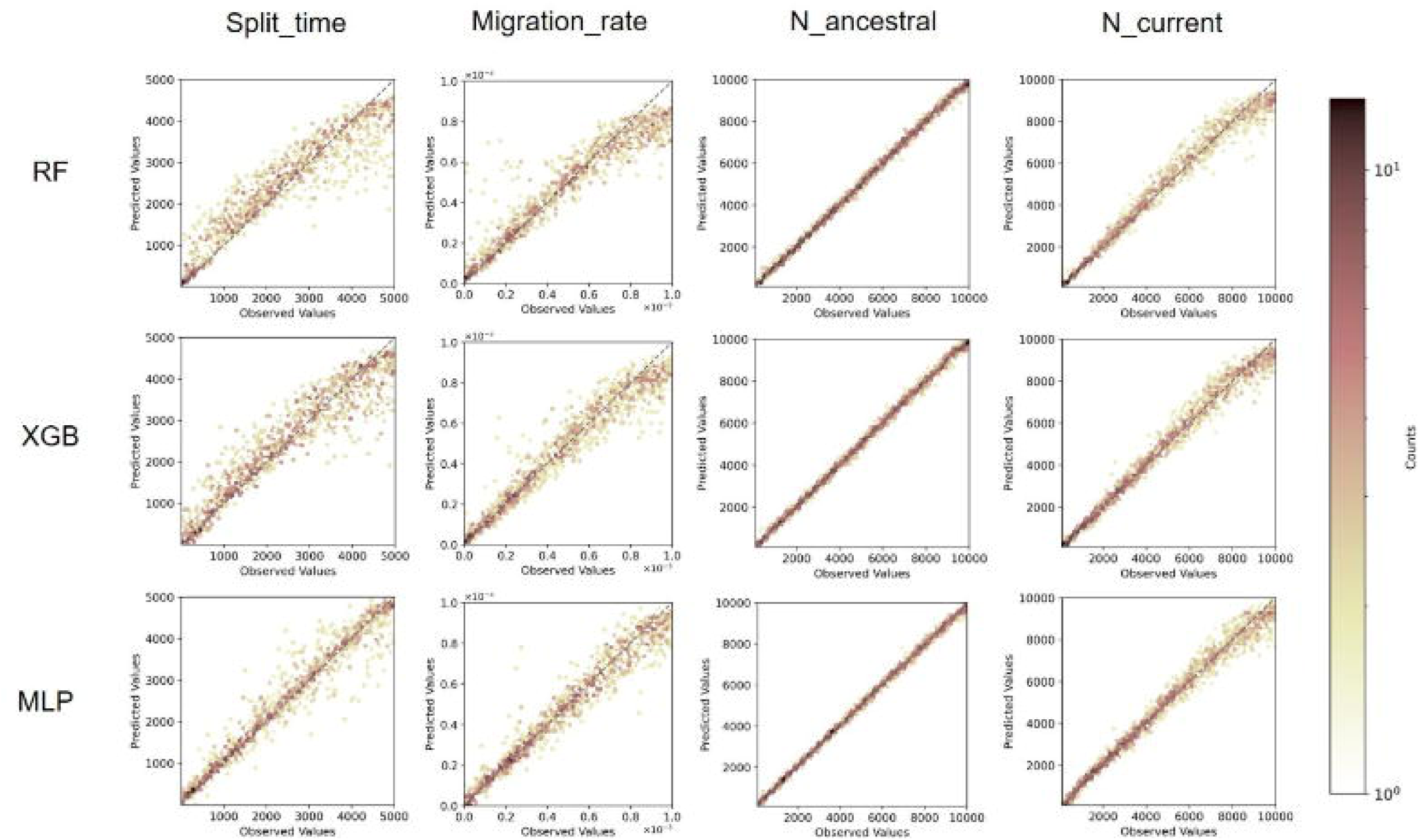
Predicted values versus observed values for the four isolation-with-migration parameters estimated with three different ML methods (RF: Random Forest, XGB: XGBoost, MLP: Multilayer perceptron).

### Comparison with ABC methods

The performance of the ABC algorithms depended on both the tolerance thresholds and the choice of using either the median or the mean of the posterior as estimator (Table S3). Whatever the tolerance threshold chosen, the three ML methods all outperformed these ABC algorithms on the same simulation set (Fig. S4). The ABC rejection algorithm always showed lower performances than the ABC loclinear and ABC neuralnet methods. Compared with the three ML methods, the performances of ABC algorithms in predicting current and ancestral effective population sizes were substantially lower. The difference between the three ML methods and the ABC algorithms was less pronounced for the migration rate parameter, but still in favour of the ML methods.

### Accuracy of the machine-learning method for parameter estimation under the SC model

We then trained the same hyper-parametrized models on the SC scenarios. In the following, *N_current* and *Growth_rate* refer to results for population 1. As the model is symmetrical and the priors are the same for both populations, the results are similar to those for population 2.

For the parameters already present in the standard IM model, MLP was again generally the best-performing method (Table 2). This was particularly noteworthy for *Split_time*, for which RF and XGB showed both a downward bias for high values of that parameter (Fig. S5). *N_ancestral* remained the most accurately predicted parameter with the lowest NMAE (4.19 x 10^-2^ for MLP). The model with the lowest RMSE for *Growth_rate* prediction was XGB (6.06 x 10^-4^), while MLP showed the lowest MAE (4.51 x 10^-4^). Finally, for the migration duration, MLP achieved the best performances in terms of both RMSE and MAE.

**Table 2.** Prediction errors on the test data set of the three ML methods, for each demographic parameter of the secondary contact with varying population sizes model. RMSE: Root Mean Square Error; MAE: Mean Absolute Error; NMAE: Mean Absolute Error normalized by the range of the parameter. The lowest errors for each parameter are indicated in bold.

| Demographic parameter | Method | RMSE | MAE (se) | NMAE (se) |
| --- | --- | --- | --- | --- |
| Split_time | RF | 8.37E+02 | 6.74E+02 (4.98E+02) | 1.37E-01 (1.02E-01) |
|  | XGB | 8.34E+02 | 6.45E+02 (5.28E+02) | 1.32E-01 (1.08E-01) |
|  | MLP | **7.32E+02** | **5.29E+02 (5.06E+02)** | **1.08E-01 (1.03E-01)** |
| Migration_rate | RF | 7.48E-04 | 5.58E-04 (4.98E-04) | 1.12E-01 (9.96E-02) |
|  | XGB | 7.33E-04 | 5.29E-04 (5.08E-04) | 1.06E-01 (1.02E-01) |
|  | MLP | **7.12E-04** | **5.08E-04 (4.99E-04)** | **1.02E-01 (9.97E-02)** |
| N_ancestral | RF | 3.29E+02 | 1.94E+02 (2.65E+02) | 4.86E-02 (6.64E-02) |
|  | XGB | 3.18E+02 | 1.88E+02 (2.57E+02) | 4.70E-02 (6.43E-02) |
|  | MLP | **3.10E+02** | **1.68E+02 (2.61E+02)** | **4.19E-02 (6.51E-02)** |
| N_current | RF | 4.72E+02 | 3.71E+02 (2.92E+02) | 9.27E-02 (7.30E-02) |
|  | XGB | **4.32E+02** | 3.34E+02 (2.75E+02) | 8.34E-02 (6.86E-02) |
|  | MLP | 4.42E+02 | **3.31E+02 (2.93E+02)** | **8.26E-02 (7.33E-02)** |
| Growth_rate | RF | 6.07E-04 | 4.89E-04 (3.60E-04) | 1.63E-01 (1.20E-01) |
|  | XGB | **6.06E-04** | 4.71E-04 (3.81E-04) | 1.57E-01 (1.27E-01) |
|  | MLP | 6.17E-04 | **4.51E-04 (4.21E-04)** | **1.50E-01 (1.40E-01)** |
| Migra-<br>tion_duration | RF | 2.19E-01 | 1.79E-01 (1.26E-01) | 1.79E-01 (1.26E-01) |
|  | XGB | 2.23E-01 | 1.78E-01 (1.35E-01) | 1.78E-01 (1.35E-01) |
|  | MLP | **2.13E-01** | **1.60E-01 (1.41E-01)** | **1.60E-01 (1.41E-01)** |

In the following sections, we focus on the standard isolation with migration model (IM).

### Assessing accuracy discrepancies across demographic scenarios

We focus in this section on MLP, as it showed the lowest MAE. We investigated if there was a heterogeneity in prediction quality according to the demographic scenarios. In order to address this question, we used regression to identify to what extent the different demographic parameters could explain prediction errors (Fig. S6). We then plotted these prediction errors as a function of the two most explanatory parameters (Fig. 3).

**Fig. 3.**
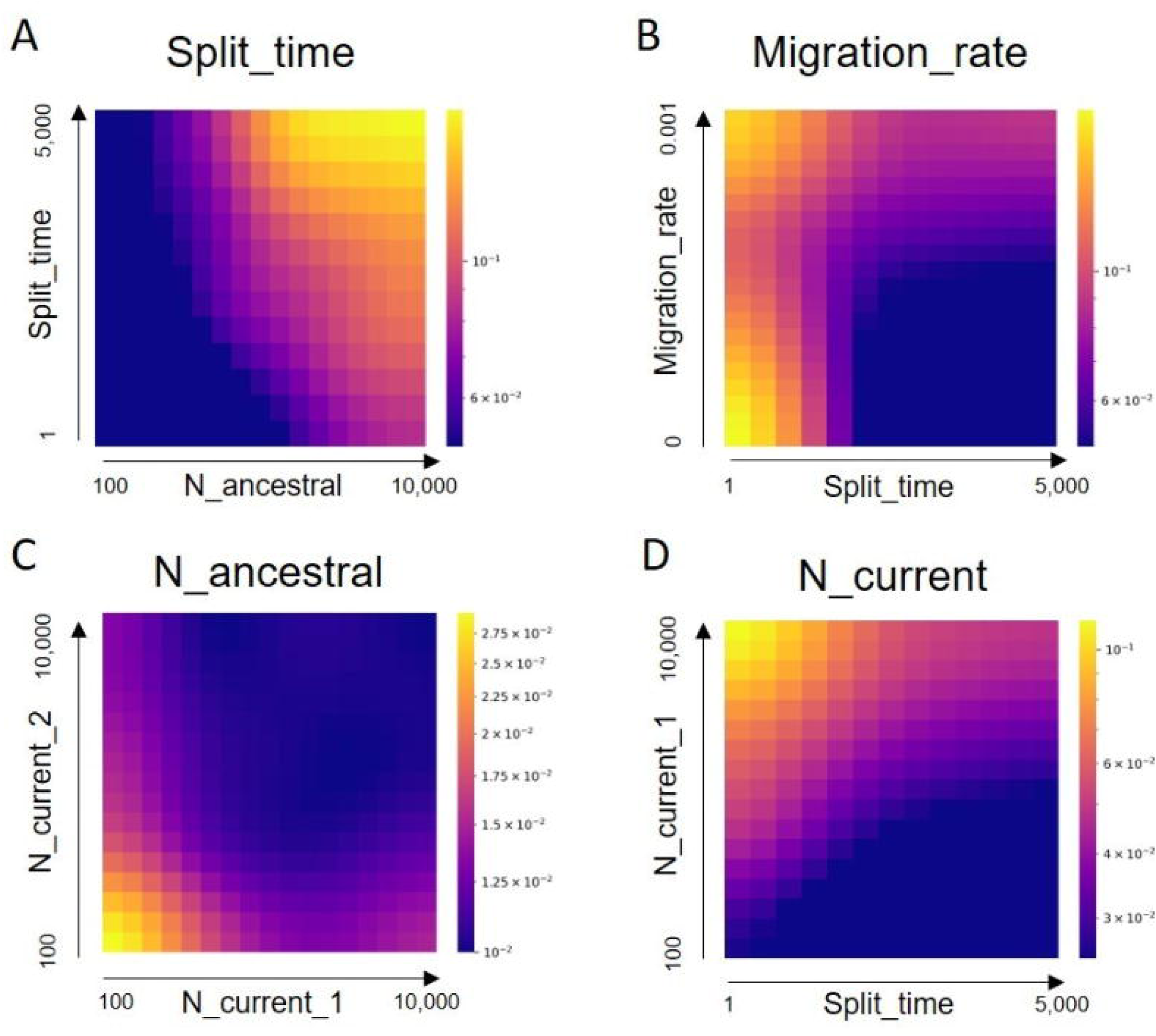
Standardized errors (ŷ_*i*_ − y_*i*_)/(y_*max*_ − y_*min*_) where y is the target demographic parameter as a function of the two variables most predictive of these errors under the MLP method. Cold colours correspond to areas of low error and therefore better model prediction, while warm colours correspond to areas of higher error.

Regarding the prediction of split time, we observed that the best performance was achieved for smaller ancestral effective population sizes (Fig. 3A). As this effective population size increased, the prediction error increased. This effect was more pronounced for older split times. When predicting the migration rate, MLP performed less efficiently for recent split times. As the split time increased, the prediction quality was higher for a lower migration rate (Fig. 3B).

As for the ancestral effective population size, the values taken by the current effective population sizes had the largest impact on the quality of the prediction. The lowest accuracy was observed for the ancestral size when both current populations showed low effective sizes. When one of the two effective population sizes was small, prediction quality remained lower than average, regardless of the effective size of the second population (Fig. 3C).

Split time was the most decisive parameter in the quality of the prediction for current effective population size parameters. As this time increased, the prediction became increasingly accurate, whereas demographic scenarios with recent split time and high current effective population sizes led to the highest errors.

### Different patterns of summary statistic usage across ML models

We estimated the importance levels of various summary statistics classes for each ML model and each demographic parameter in order to determine to which extent they differ in that aspect. We computed the importance of each variable, using the permutation feature importance (PFI) method for the three ML methods, which computes the extent to which the quality of a prediction of a given model decreases when a single variable is randomly shuffled.

We observed first that RF and XGB allocated quite similar importance levels to the various classes of summary statistics (Fig. 4), while the MLP method generally showed different behaviours. For *Split_time*, the *SFS* class stood out for the two ensemble methods, with importance levels of 0.71 and 0.66 respectively. The *JSFS* class was second in terms of importance (importance levels of 0.10 and 0.16 for RF and XGB, respectively), but the gap was high with the *SFS* class. Conversely, for MLP, the *JSFS* class was the most important, with an importance of 0.63, ahead of the *SFS* and *AFIBS* classes. The differences among methods were much less pronounced for the *Migration_rate* parameter, where the *AFIBS*, *SFS*, and foremost *JSFS* classes were used predominantly by all methods. Nevertheless, the XGB and RF models also gave significant importance to *Fst*, unlike the MLP model.

**Fig. 4.**
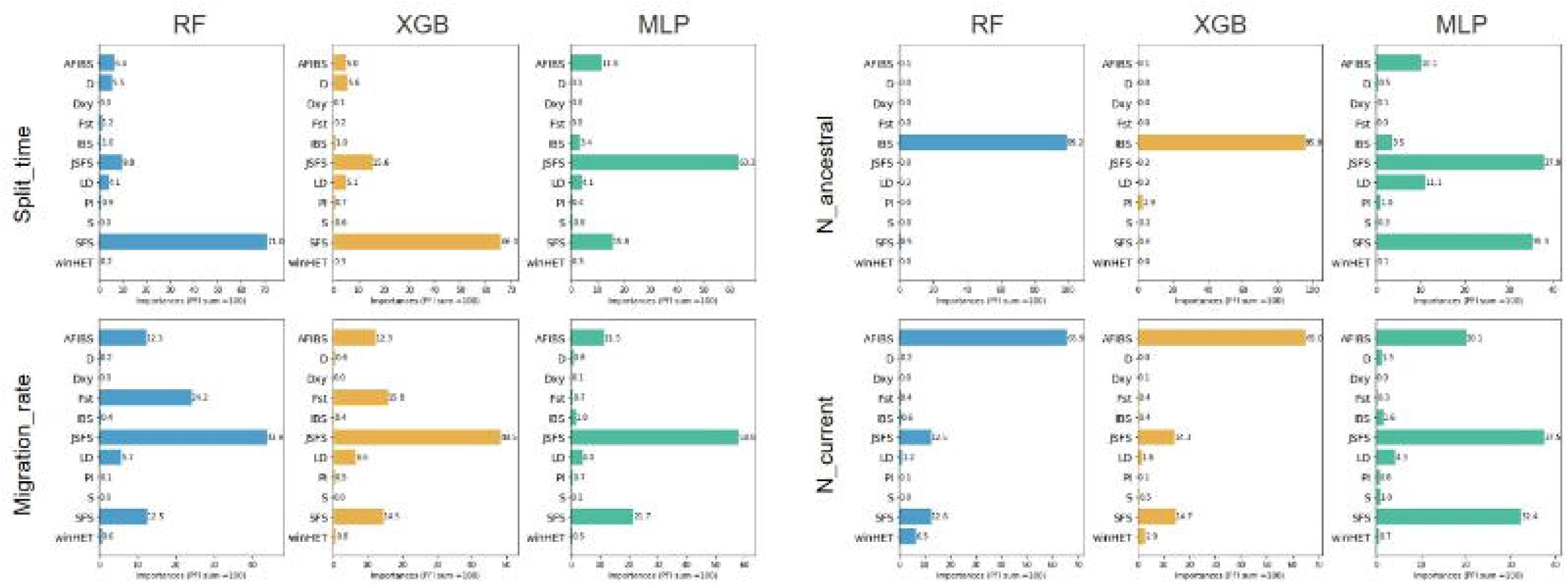
Importance levels of the different classes of summary statistics as a percentage of the total for each demographic parameter and each method (RF, XGB, MLP).

The differences were much more striking for the effective population sizes. For the ancestral effective population size, RF and XGB focused almost exclusively on the *IBS* class, with importance levels of 0.99 and 0.96, respectively. Conversely, MLP used the different classes much more evenly, with five classes showing noticeable levels of importance. The discrepancy among methods was a bit lower but still striking for the current effective population sizes. All methods used several classes of summary statistics, but the importance levels were strongly unbalanced toward *AFIBS* for RF and XGB, with values 0.66 and 0.65 respectively, while MLP used again the different classes more evenly.

This similarity in the variables used by XGB and RF was confirmed by observing the individual importance levels of each summary statistics (see Fig. S7). In the case of split time, the variables related to *SFS* were of the utmost importance for the predictions made by these two methods. In a similar manner, these methods assigned the greatest importance to *Fst*, followed by statistics from the *JSFS* class for predicting the migration rate. The most important variables for predicting the ancestral effective population size belonged all to the *IBS* class, while for the current effective population size these variables belonged to different classes: *AFIBS*, *JSFS* and *winHET*. The most important summary statistics chosen by MLP differed strikingly from those chosen by RF and XGB. They also exhibited markedly lower importance values. Indeed, the maximum importance attributed to a single feature by MLP was 2.6% for the *Migration_rate* parameter. For this method, importance was clearly spread over a much larger number of variables than for the two other methods (Fig. S8).

### Towards improved explainability

The ML methods do not directly provide the impact of each individual statistic on the parameter estimation, due to their inherent complexity, non-linear relationships, or the fact that they are a combination of a set of predictors. This makes the interpretation of the individual impact of each summary statistics more complex. While some methods such as the PFI method used above are effective in revealing features that are important for prediction, they reach their limit when it comes to quantifying the impact of features while taking into account how they interact with each other. Interpretation techniques may then prove necessary to provide insights into the role of summary statistics in complex models. We therefore used SHAP (Lundberg & Lee, 2017), a method based on Shapley values that measures the average marginal contribution of each feature when combined with all other to make a prediction for a given parameter. In addition to identifying the most important summary statistics for each parameter, this method also allows investigating finely how these statistics affect the model outputs, quantifying both the direction and magnitude of their contributions to the prediction of each parameter. We illustrate the contributions of this method in the IM model for the *Split_time* predictions obtained with the XGB model, which combined high accuracy with an ability to bring out statistics with substantial importance (Fig. S7).

Focusing first on the summary statistics with the greatest impact, we observed that the frequency of tripletons in the overall population (*SFS_3_all-mu*) showed the highest mean of absolute SHAP values (Fig. 5A). This statistic was also the one that showed the highest importance (Fig. S7). The distribution of SHAP values for the most important variables allowed us to understand the impact of individual summary statistics on predictions (Fig. 5B). We observe that *SFS_3_all-mu* had the greatest upward and downward impact on the prediction of *Split_time*. While the highest values of this variable almost always had an upward impact, we see that intermediate values had a substantial impact, particularly downward, when combined with other statistics (Fig. 5C). We also observed that for the variable *SFS_19_p1-mu*, which represents the frequency of SNPs with the ancestral allele found in a single haplotype, higher values have a greater impact on the parameter compared to lower values, which have a more limited impact (Fig. 5B).

**Fig. 5.**
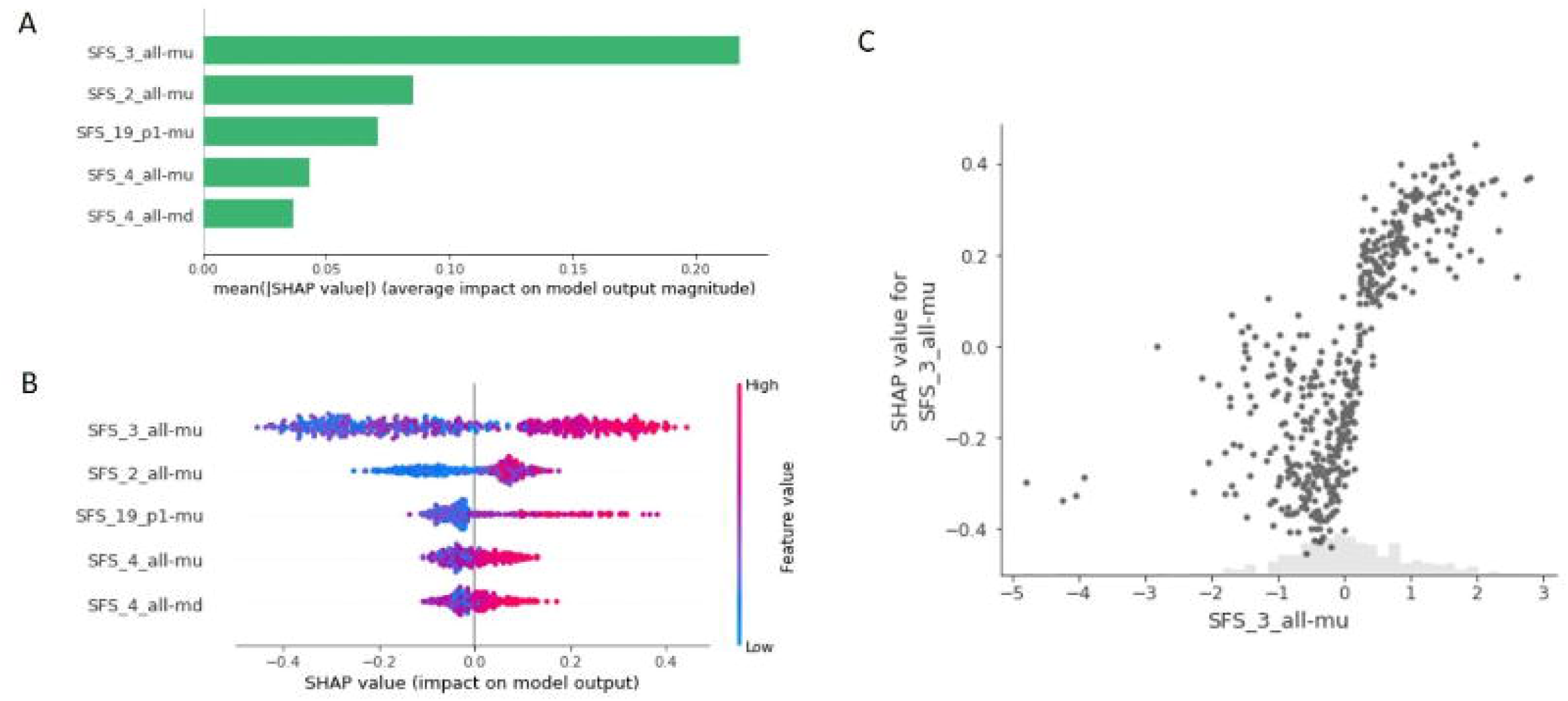
(A) Top features with the highest means of absolute SHAP values for the XGB model on the *Split_time* parameter. (B) Feature effects indicated by the Shapley value on the x-axis for the top 5 features with their values represented by colour. Each point corresponds to one instance. (C) Shapley values as a function of the normalized *SFS_3_all-mu* (grey points), distribution of the normalized *SFS_3_all-mu* (grey bars).

## Discussion

We have shown here how an approach combining a large number of summary statistics and machine learning methods was particularly effective in predicting demographic parameters in scenarios involving two connected populations. These methods do not require preselecting a subset of summary statistics in order to reduce the number of variables. Both ensemble and neural network methods are indeed suitable for learning from a very large number of statistics, some of which being highly correlated, and some being aggregated indicators that can be drawn from others. We showed that MLP overall outperformed the RF and XGB methods in parameter prediction for the considered IM and SC model. Other demographic and genetic modelling assumptions could be tested in future works using the available scripts, as well as the effect of the number of simulations on accuracy. Other prior distributions could also be tested, such as log-uniform distributions, which are of particular interest when we are interested in inferring the order of magnitude of a parameter. Although achieving lower accuracy, RF and XGB still showed good performances. Depending on the metric and degree of accuracy required, they could still be considered in some cases, as they are less computer-intensive.

The intricate effects of the demographic parameters on the summary statistics undermine the predictive power of these statistics. We therefore emphasize the importance of studying the impact of confounding factors (here, the other demographic parameters) and the heterogeneity of prediction accuracy across the parameter space. Within the isolation with migration framework, we identified which demographic parameters were most decisive in the accuracy of the prediction of our target parameter, and visualized how this accuracy varies as a function of these most explanatory parameters. Taking such an approach when building a new model for a given evolutionary question ensures a precise picture of the inference strengths and weaknesses.

The three machine learning methods showed contrasting patterns in terms of summary statistics usage for predicting demographic parameters in the modelling settings of the IM model. Although XGB showed generally better performances than RF, both methods used quite similar learning schemes. MLP, on the other hand, used strikingly different classes of summary statistics than the two other methods. Furthermore, whereas RF and XGB focused on a limited number of summary statistics to which they gave a high importance, MLP used a much larger number of statistics, each being given a small importance. In other words, the information used by MLP was spread much more evenly over the different statistics. Based on a higher degree of complexity, this method therefore showed its higher capacity to capture intricate patterns and complex relationships from a high number of variables, as for instance those of the *JSFS* class.

Conversely, by pointing at a limited number of important statistics for prediction, tree-based methods reveal few important summary statistics for inference. We therefore note that an error on one of the most important summary statistics is likely to have a strong impact on the estimation of the demographic parameter. However, this has the advantage of allowing us to identify and select the specific classes of summary statistics to which we may wish to restrict ourselves for a particular demographic model, enhancing the interpretability of the results. For this purpose, we can rely on the recently developed interpretability method SHAP (Lundberg & Lee, 2017), which was notably used to interpret mutation rate predictions from a neural network based on SFS (Burger et al., 2022) and in a deep learning framework for selection inference (Cecil & Sugden, 2023). This method allowed us to assess how the values of a given summary statistic affect the prediction of the demographic parameters under study. Ultimately, while the methods developed here were tested in specific demographic scenarios, with mutation and recombination rates consistent with known values for humans, they could be adapted to other contexts, in which their performances could be tested with the same procedure as we used here. The impact of the sample size and of the number of loci could also be studied either using realistic ranges or matching the exact values of the user’s dataset.

Note also that the methods that we implemented here are likely to undergo several developments in the future. Our simulation approach was based indeed on a strictly neutral model, without any selective pressure. However, it has been shown that positive selection (Schrider et al., 2016), as well as background and purifying selection (Ewing & Jensen, 2016; Pouyet et al., 2018; Johri et al., 2021), have an impact on demographic inferences. Therefore, while it is difficult for other approaches such as composite-likelihood and SMC-based methods to account for selection explicitly, methods developed in this study could naturally be extended to models that incorporate these selective effects, allowing for a more comprehensive assessment of their impact on demographic inferences.

Similarly, we simulated sequences with even mutation and recombination rates, and without sequencing errors. A relaxation of these assumptions could also be explored (see e.g. Boitard et al., 2016; Jay et al., 2019). We further point out that while all demographic inference methods are sensitive to model assumptions, simulation-based approaches including ABC methods are also sensitive to simulation misspecification. Since the prediction of demographic parameters is restricted to predefined prior bounds, it is essential to ensure that these bounds are set carefully, without being too restrictive. An additional area of focus could be to investigate how these methods perform for model selection, often a preliminary step to parameter estimation.

Finally, even if we showed here the high predictive power of our methods, we suggest to compare their efficiency with that of methods that rely on the direct application of deep learning methods to raw genomic data, in order to infer demographic (Sanchez et al., 2021) or selection events (Flagel et al., 2019; Torada et al., 2019). By learning more complex patterns from the data that may not be captured by summary statistics devised beforehand, these methods may prove particularly effective in case of complex demographic models. However, as they do not benefit from the guidance provided by the summary statistics, it is necessary to compare their performances with those of methods such as the ones that we developed here, as carried out by Sanchez et al. (2021) for the estimation of the effective size of a population over time. In this context, they showed that current neural networks yielded accurate predictions without requiring handcrafted features, but summary statistics were already compelling if carefully optimizing the inference model.

## Data availability

Scripts used to generate simulated data and run the demographic inferences are available on github at: https://github.com/amquelin/SML_demographic_inference.

## Acknowledgements

We thank Emilia Huerta-Sanchez and Camille Roux for insightful discussions. We thank the editor and three anonymous reviewers for their comments on the manuscript.

## Authors’ contributions

All authors conceived and designed the study. AQ implemented the approach. All authors analysed the results. AQ wrote the manuscript draft. FA and FJ reviewed and edited the draft. All authors read and approved the final manuscript.

## References

Abascal, F., Corvelo, A., Cruz, F., Villanueva-Cañas, J. L., Vlasova, A., Marcet-Houben, M. et al. (2016). Extreme genomic erosion after recurrent demographic bottlenecks in the highly endangered Iberian lynx. Genome Biology, 17(1), 251. 10.1186/s13059-016-1090-1

Agarap, A. F. (2019). Deep Learning using Rectified Linear Units (ReLU) (arXiv:1803.08375). arXiv. 10.48550/arXiv.1803.08375

Akey, J. M., Eberle, M. A., Rieder, M. J., Carlson, C. S., Shriver, M. D., Nickerson, D. A. et al. (2004). Population History and Natural Selection Shape Patterns of Genetic Variation in 132 Genes. PLOS Biology, 2(10), e286. 10.1371/journal.pbio.0020286

Bai, W.-N., Yan, P.-C., Zhang, B.-W., Woeste, K. E., Lin, K., & Zhang, D.-Y. (2018). Demographically idiosyncratic responses to climate change and rapid Pleistocene diversification of the walnut genus Juglans (Juglandaceae) revealed by whole-genome sequences. New Phytologist, 217(4), 1726– 1736. 10.1111/nph.14917

Baumdicker, F., Bisschop, G., Goldstein, D., Gower, G., Ragsdale, A. P., Tsambos, G. et al. (2021). Efficient ancestry and mutation simulation with msprime 1.0. Genetics, 220(3), iyab229. 10.1093/genetics/iyab229

Beaumont, M. A. (2010). Approximate Bayesian Computation in Evolution and Ecology. Annual Review of Ecology, Evolution, and Systematics, 41(1), 379–406. 10.1146/annurev-ecolsys-102209-144621

Beaumont, M. A., Zhang, W., & Balding, D. J. (2002). Approximate Bayesian computation in population genetics. Genetics, 162(4), 2025–2035. https://www.ncbi.nlm.nih.gov/pmc/articles/PMC1462356/

Beichman, A. C., Huerta-Sanchez, E., & Lohmueller, K. E. (2018). Using Genomic Data to Infer Historic Population Dynamics of Nonmodel Organisms. Annual Review of Ecology, Evolution, and Systematics, 49(1), 433–456. 10.1146/annurev-ecolsys-110617-062431

Bertorelle, G., Benazzo, A., & Mona, S. (2010). ABC as a flexible framework to estimate demography over space and time: Some cons, many pros. Molecular Ecology, 19(13), 2609–2625. 10.1111/j.1365-294X.2010.04690.x

Blum, M. G. B., & Francois, O. (2010). Non-linear regression models for Approximate Bayesian Computation. Statistics and Computing, 20(1), 63–73. 10.1007/s11222-009-9116-0

Boitard, S., Rodríguez, W., Jay, F., Mona, S., & Austerlitz, F. (2016). Inferring Population Size History from Large Samples of Genome-Wide Molecular Data—An Approximate Bayesian Computation Approach. PLOS Genetics, 12(3), e1005877. 10.1371/journal.pgen.1005877

Breiman, L. (2001). Random Forests. Machine Learning, 45(1), 5–32. 10.1023/A:1010933404324

Browning, S. R., & Browning, B. L. (2015). Accurate Non-parametric Estimation of Recent Effective Population Size from Segments of Identity by Descent. American Journal of Human Genetics, 97(3), 404–418. 10.1016/j.ajhg.2015.07.012

Burger, K. E., Pfaffelhuber, P., & Baumdicker, F. (2022). Neural networks for self-adjusting mutation rate estimation when the recombination rate is unknown. PLOS Computational Biology, 18(8), e1010407. 10.1371/journal.pcbi.1010407

Campbell, C. D., Chong, J. X., Malig, M., Ko, A., Dumont, B. L., Han, L., Vives, L., O’Roak, B. J. et al. (2012). Estimating the human mutation rate using autozygosity in a founder population. Nature Genetics, 44(11), 1277– 1281. 10.1038/ng.2418

Cavalli-Sforza, L. L., Menozzi, P., & Piazza, A. (1994). The history and geography of human genes. Princeton university press.

Cecil, R. M., & Sugden, L. A. (2023). On convolutional neural networks for selection inference: Revealing the effect of preprocessing on model learning and the capacity to discover novel patterns. PLOS Computational Biology, 19(11), e1010979. 10.1371/journal.pcbi.1010979

Chai, T., & Draxler, R. R. (2014). Root mean square error (RMSE) or mean absolute error (MAE)? – Arguments against avoiding RMSE in the literature. Geoscientific Model Development, 7(3), 1247–1250. 10.5194/gmd-7-1247-2014

Chen, T., & Guestrin, C. (2016). XGBoost: A Scalable Tree Boosting System. Proceedings of the 22nd ACM SIGKDD International Conference on Knowledge Discovery and Data Mining, 785–794. 10.1145/2939672.2939785

Chikhi, L., Rodríguez, W., Grusea, S., Santos, P., Boitard, S., & Mazet, O. (2018). The IICR (inverse instantaneous coalescence rate) as a summary of genomic diversity: Insights into demographic inference and model choice. Heredity, 120(1), 13–24.

Collin, F., Durif, G., Raynal, L., Lombaert, E., Gautier, M., Vitalis, R. et al. (2021). Extending approximate Bayesian computation with supervised machine learning to infer demographic history from genetic polymorphisms using DIYABC Random Forest. Molecular Ecology Resources, 21(8), 2598–2613. 10.1111/1755-0998.13413

Csilléry, K., François, O., & Blum, M. G. B. (2012). abc: An R package for approximate Bayesian computation (ABC). Methods in Ecology and Evolution, 3(3), 475–479. 10.1111/j.2041-210X.2011.00179.x

Der Sarkissian, C., Ermini, L., Schubert, M., Yang, M. A., Librado, P., Fumagalli, M., et al. (2015). Evolutionary Genomics and Conservation of the Endangered Przewalski’s Horse. Current Biology, 25(19), 2577–2583. 10.1016/j.cub.2015.08.032

Dong, F., Kuo, H.-C., Chen, G.-L., Wu, F., Shan, P.-F., Wang, J. et al. (2021). Population genomic, climatic and anthropogenic evidence suggest the role of human forces in endangerment of green peafowl (Pavo muticus). Proceedings of the Royal Society B: Biological Sciences, 288(1948), 20210073. 10.1098/rspb.2021.0073

Estoup, A., Lombaert, E., Marin, J.-M., Guillemaud, T., Pudlo, P., Robert, C. P. et al. (2012). Estimation of demo-genetic model probabilities with Approximate Bayesian Computation using linear discriminant analysis on summary statistics. Molecular Ecology Resources, 12(5), 846–855. 10.1111/j.1755-0998.2012.03153.x

Ewing, G. B., & Jensen, J. D. (2016). The consequences of not accounting for background selection in demographic inference. Molecular Ecology, 25(1), 135–141. 10.1111/mec.13390

Fedorov, V. B., Trucchi, E., Goropashnaya, A. V., Waltari, E., Whidden, S. E., & Stenseth, N. Chr. (2020). Impact of past climate warming on genomic diversity and demographic history of collared lemmings across the Eurasian Arctic. Proceedings of the National Academy of Sciences, 117(6), 3026–3033. 10.1073/pnas.1913596117

Flagel, L., Brandvain, Y., & Schrider, D. R. (2019). The Unreasonable Effectiveness of Convolutional Neural Networks in Population Genetic Inference. Molecular Biology and Evolution, 36(2), 220–238. 10.1093/molbev/msy224

Ghirotto, S., Vizzari, M. T., Tassi, F., Barbujani, G., & Benazzo, A. (2021). Distinguishing among complex evolutionary models using unphased whole-genome data through random forest approximate Bayesian computation. Molecular Ecology Resources, 21(8), 2614–2628. 10.1111/1755-0998.13263

Gutenkunst, R. N., Hernandez, R. D., Williamson, S. H., & Bustamante, C. D. (2009). Inferring the Joint Demographic History of Multiple Populations from Multidimensional SNP Frequency Data. PLOS Genetics, 5(10), e1000695. 10.1371/journal.pgen.1000695

Harris, K., & Nielsen, R. (2013). Inferring Demographic History from a Spectrum of Shared Haplotype Lengths. PLOS Genetics, 9(6), e1003521. 10.1371/journal.pgen.1003521

Haykin, S. (1994). Neural networks: A comprehensive foundation. Prentice Hall PTR.

Henn, B. M., Cavalli-Sforza, L. L., & Feldman, M. W. (2012). The great human expansion. Proceedings of the National Academy of Sciences of the United States of America, 109(44), 17758–17764. 10.1073/pnas.1212380109

Jabot, F., & Lohier, T. (2016). Non-random correlation of species dynamics in tropical tree communities. Oikos, 125(12), 1733–1742. 10.1111/oik.03103

Jay, F., Boitard, S., & Austerlitz, F. (2019). An ABC Method for Whole-Genome Sequence Data: Inferring Paleolithic and Neolithic Human Expansions. Molecular Biology and Evolution, 36(7), 1565–1579. 10.1093/molbev/msz038

Johri, P., Pfeifer, S. P., & Jensen, J. D. (2023). Developing an Evolutionary Baseline Model for Humans: Jointly Inferring Purifying Selection with Population History. Molecular Biology and Evolution, 40(5), msad100. 10.1093/molbev/msad100

Johri, P., Riall, K., Becher, H., Excoffier, L., Charlesworth, B., & Jensen, J. D. (2021). The Impact of Purifying and Background Selection on the Inference of Population History: Problems and Prospects. Molecular Biology and Evolution, 38(7), 2986–3003. 10.1093/molbev/msab050

Kelleher, J., Etheridge, A. M., & McVean, G. (2016). Efficient Coalescent Simulation and Genealogical Analysis for Large Sample Sizes. PLOS Computational Biology, 12(5), e1004842. 10.1371/journal.pcbi.1004842

Kong, A., Frigge, M. L., Masson, G., Besenbacher, S., Sulem, P., Magnusson, G. et al. (2012). Rate of de novo mutations and the importance of father’s age to disease risk. Nature, 488(7412), Article 7412. 10.1038/nature11396

Korfmann, K., Gaggiotti, O. E., & Fumagalli, M. (2023). Deep learning in population genetics. Genome Biology and Evolution, evad008. 10.1093/gbe/evad008

Li, H., & Durbin, R. (2011). Inference of human population history from individual whole-genome sequences. Nature, 475(7357), Article 7357. 10.1038/nature10231

Liepe, J., Kirk, P., Filippi, S., Toni, T., Barnes, C. P., & Stumpf, M. P. H. (2014). A framework for parameter estimation and model selection from experimental data in systems biology using approximate Bayesian computation. Nature Protocols, 9(2), Article 2. 10.1038/nprot.2014.025

Liu, X., & Fu, Y.-X. (2015). Exploring Population Size Changes Using SNP Frequency Spectra. Nature Genetics, 47(5), 555–559. 10.1038/ng.3254

Lohmueller, K. E. (2014). The Impact of Population Demography and Selection on the Genetic Architecture of Complex Traits. PLOS Genetics, 10(5), e1004379. 10.1371/journal.pgen.1004379

Lundberg, S. M., & Lee, S.-I. (2017). A Unified Approach to Interpreting Model Predictions. Advances in Neural Information Processing Systems, 30. https://papers.nips.cc/paper_files/paper/2017/hash/8a20a8621978632d76c43dfd28b67767-Abstract.html

Marchi, N., Schlichta, F., & Excoffier, L. (2021). Demographic inference. Current Biology, 31(6), R276–R279. 10.1016/j.cub.2021.01.053

Marjoram, P., & Wall, J. D. (2006). Fast ‘coalescent’ simulation. BMC Genetics, 7(1), 16. 10.1186/1471-2156-7-16

McKinley, T. J., Vernon, I., Andrianakis, I., McCreesh, N., Oakley, J. E., Nsubuga, R. et al. (2018). Approximate Bayesian Computation and Simulation-Based Inference for Complex Stochastic Epidemic Models. Statistical Science, 33(1), 4–18. 10.1214/17-STS618

McVean, G. A. T., & Cardin, N. J. (2005). Approximating the coalescent with recombination. Philosophical Transactions of the Royal Society B: Biological Sciences, 360(1459), 1387–1393. 10.1098/rstb.2005.1673

Mondal, M., Bertranpetit, J., & Lao, O. (2019). Approximate Bayesian computation with deep learning supports a third archaic introgression in Asia and Oceania. Nature Communications, 10(1), Article 1. 10.1038/s41467-018-08089-7

Nei, M., & Li, W. H. (1979). Mathematical model for studying genetic variation in terms of restriction endonucleases. Proceedings of the National Academy of Sciences, 76(10), 5269–5273. 10.1073/pnas.76.10.5269

Palamara, P. F., Lencz, T., Darvasi, A., & Pe’er, I. (2012). Length Distributions of Identity by Descent Reveal Fine-Scale Demographic History. American Journal of Human Genetics, 91(5), 809–822. 10.1016/j.ajhg.2012.08.030

Pedregosa, F., Varoquaux, G., Gramfort, A., Michel, V., Thirion, B., Grisel, O. et al. (2011). Scikit-learn: Machine Learning in Python. Journal of Machine Learning Research, 12(85), 2825–2830. http://jmlr.org/papers/v12/pedregosa11a.html

Pouyet, F., Aeschbacher, S., Thiéry, A., & Excoffier, L. (2018). Background selection and biased gene conversion affect more than 95% of the human genome and bias demographic inferences. Elife, 7, e36317.

Pritchard, J. K., Seielstad, M. T., Perez-Lezaun, A., & Feldman, M. W. (1999). Population growth of human Y chromosomes: A study of Y chromosome microsatellites. Molecular Biology and Evolution, 16(12), 1791–1798. 10.1093/oxfordjournals.molbev.a026091

Pudlo, P., Marin, J.-M., Estoup, A., Cornuet, J.-M., Gautier, M., & Robert, C. P. (2016). Reliable ABC model choice via random forests. Bioinformatics, 32(6), 859–866. 10.1093/bioinformatics/btv684

Pujolar, J. M., Dalén, L., Hansen, M. M., & Madsen, J. (2017). Demographic inference from whole-genome and RAD sequencing data suggests alternating human impacts on goose populations since the last ice age. Molecular Ecology, 26(22), 6270–6283. 10.1111/mec.14374

Raynal, L., Marin, J.-M., Pudlo, P., Ribatet, M., Robert, C. P., & Estoup, A. (2019). ABC random forests for Bayesian parameter inference. Bioinformatics, 35(10), 1720–1728. 10.1093/bioinformatics/bty867

Sanchez, T., Cury, J., Charpiat, G., & Jay, F. (2021). Deep learning for population size history inference: Design, comparison and combination with approximate Bayesian computation. Molecular Ecology Resources, 21(8), 2645–2660.

Scally, A., & Durbin, R. (2012). Revising the human mutation rate: Implications for understanding human evolution. Nature Reviews Genetics, 13(10), Article 10. 10.1038/nrg3295

Schiffels, S., & Durbin, R. (2014). Inferring human population size and separation history from multiple genome sequences. Nature Genetics, 46(8), Article 8. 10.1038/ng.3015

Schraiber, J. G., & Akey, J. M. (2015). Methods and models for unravelling human evolutionary history. Nature Reviews. Genetics, 16(12), 727–740. 10.1038/nrg4005

Schrider, D. R., & Kern, A. D. (2018). Supervised Machine Learning for Population Genetics: A New Paradigm. Trends in Genetics!Z: TIG, 34(4), 301–312. 10.1016/j.tig.2017.12.005

Schrider, D. R., Shanku, A. G., & Kern, A. D. (2016). Effects of linked selective sweeps on demographic inference and model selection. Genetics, 204(3), 1207–1223.

Shapley, L. S. (1953). 17. A Value for n-Person Games. In H. W. Kuhn & A. W. Tucker (Eds.), Contributions to the Theory of Games (AM-28), Volume II (pp. 307–318). Princeton University Press. 10.1515/9781400881970-018

Sheehan, S., Harris, K., & Song, Y. S. (2013). Estimating Variable Effective Population Sizes from Multiple Genomes: A Sequentially Markov Conditional Sampling Distribution Approach. Genetics, 194(3), 647–662. 10.1534/genetics.112.149096

Sheehan, S., & Song, Y. S. (2016). Deep Learning for Population Genetic Inference. PLOS Computational Biology, 12(3), e1004845. 10.1371/journal.pcbi.1004845

Tajima, F. (1989). Statistical method for testing the neutral mutation hypothesis by DNA polymorphism. Genetics, 123(3), 585–595. 10.1093/genetics/123.3.585

Tanaka, M. M., Francis, A. R., Luciani, F., & Sisson, S. A. (2006). Using Approximate Bayesian Computation to Estimate Tuberculosis Transmission Parameters From Genotype Data. Genetics, 173(3), 1511–1520. 10.1534/genetics.106.055574

Tavaré, S., Balding, D. J., Griffiths, R. C., & Donnelly, P. (1997). Inferring Coalescence Times from DNA Sequence Data. Genetics, 145(2), 505–518. https://www.ncbi.nlm.nih.gov/pmc/articles/PMC1207814/

Terhorst, J., Kamm, J. A., & Song, Y. S. (2017). Robust and scalable inference of population history from hundreds of unphased whole genomes. Nature Genetics, 49(2), 303–309. 10.1038/ng.3748

Theunert, C., Tang, K., Lachmann, M., Hu, S., & Stoneking, M. (2012). Inferring the History of Population Size Change from Genome-Wide SNP Data. Molecular Biology and Evolution, 29(12), 3653–3667. 10.1093/molbev/mss175

Thouzeau, V., Affholder, A., Mennecier, P., Verdu, P., & Austerlitz, F. (2022). Inferring linguistic transmission between generations at the scale of individuals. Journal of Language Evolution, 7(2), 200–212. 10.1093/jole/lzac009

Thouzeau, V., Mennecier, P., Verdu, P., & Austerlitz, F. (2017). Genetic and linguistic histories in Central Asia inferred using approximate Bayesian computations. Proceedings of the Royal Society B: Biological Sciences, 284(1861), 20170706. 10.1098/rspb.2017.0706

Toni, T., Welch, D., Strelkowa, N., Ipsen, A., & Stumpf, M. P. H. (2008). Approximate Bayesian computation scheme for parameter inference and model selection in dynamical systems. Journal of The Royal Society Interface, 6(31), 187–202. 10.1098/rsif.2008.0172

Torada, L., Lorenzon, L., Beddis, A., Isildak, U., Pattini, L., Mathieson, S. et al. (2019). ImaGene: A convolutional neural network to quantify natural selection from genomic data. BMC Bioinformatics, 20(Suppl 9), 337. 10.1186/s12859-019-2927-x

Wakeley, J., & Hey, J. (1997). Estimating Ancestral Population Parameters. Genetics, 145(3), 847–855. https://10.1093/genetics/145.3.847

Wood, M. F., Simon N. (2018). ABC in Ecological Modelling. In Handbook of Approximate Bayesian Computation. Chapman and Hall/CRC.

Wright, S. (1950). Genetical Structure of Populations. Nature, 166(4215), 247–249. 10.1038/166247a0

