## Supplementary_information for "Assessing simulation-based supervised machine learning for demographic parameter inference from genomic data"

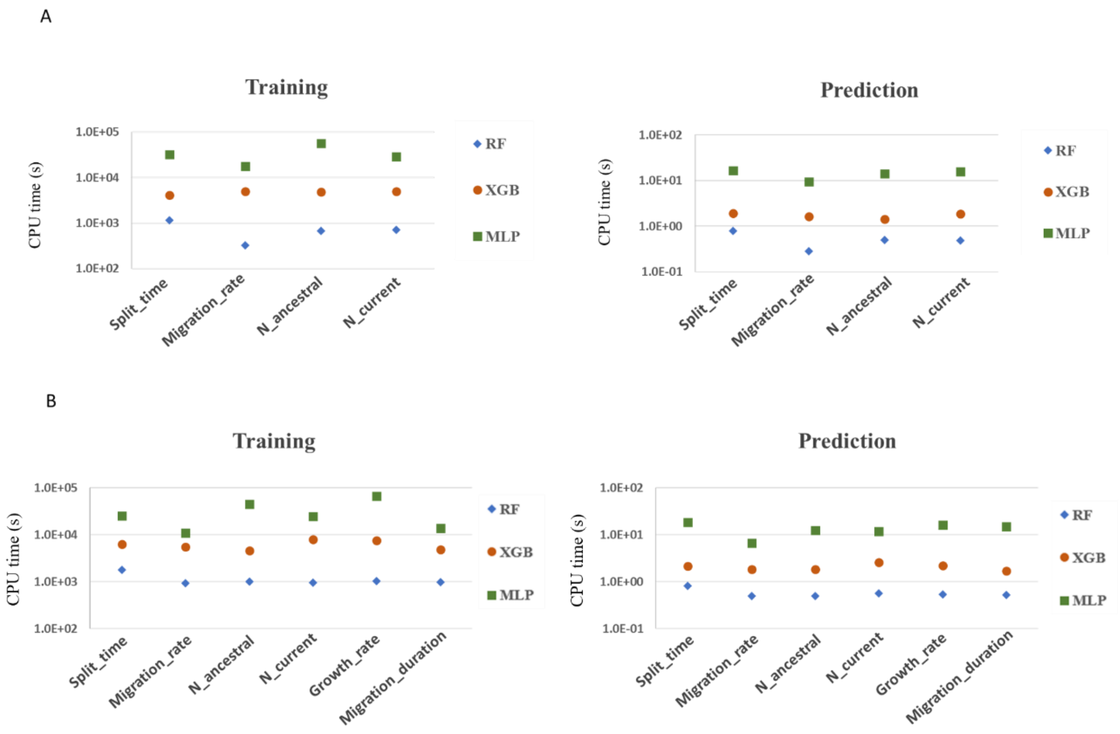


Figure S1. Comparison of the CPU time expressed as the number of seconds necessary to perform training for the various demographic parameters on the 5,000 scenarios in the training set, and their prediction on the 2,500 scenarios in the test set. A – Isolation with migration model. B – Secondary contact with varying population sizes model.

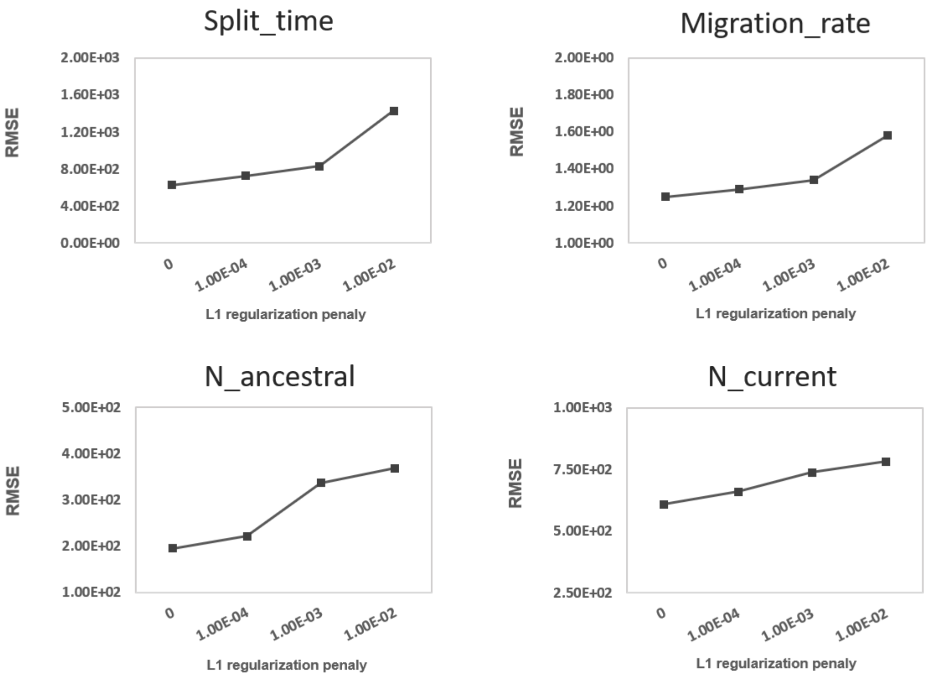


Figure S2. Evolution of RMSE (computed on the validation set) for MLP models while introducing different levels of L1 regularization penalty (x-axis).

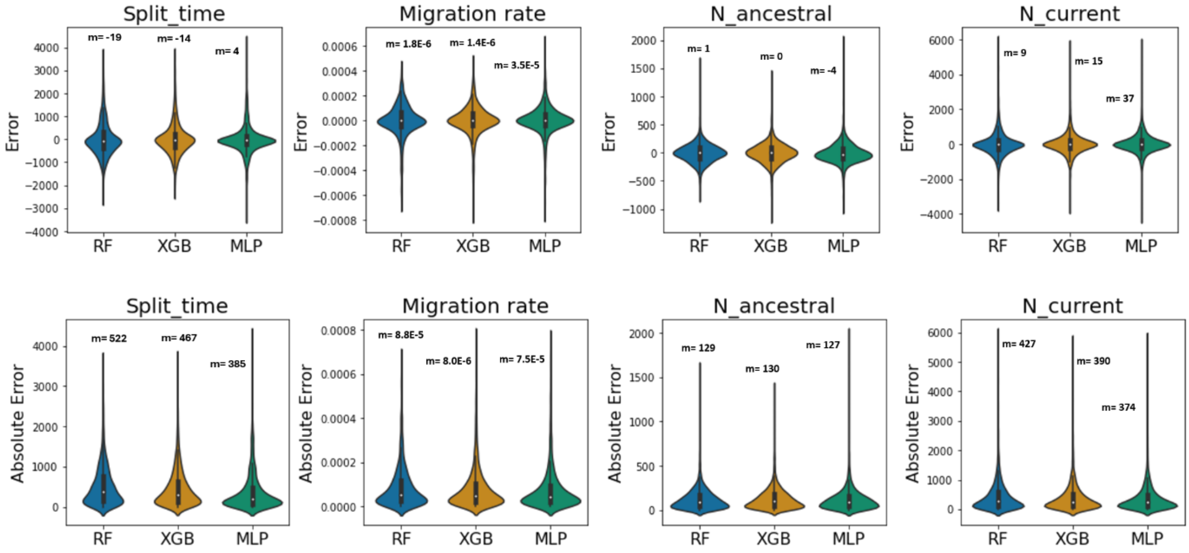


Figure S3. Distributions of the errors (top) and absolute errors (bottom) for the demographic parameters evaluated with the three different machine learning methods. We indicate by *m* the mean values of these distributions. They correspond to the bias for errors and the MAE for absolute errors.

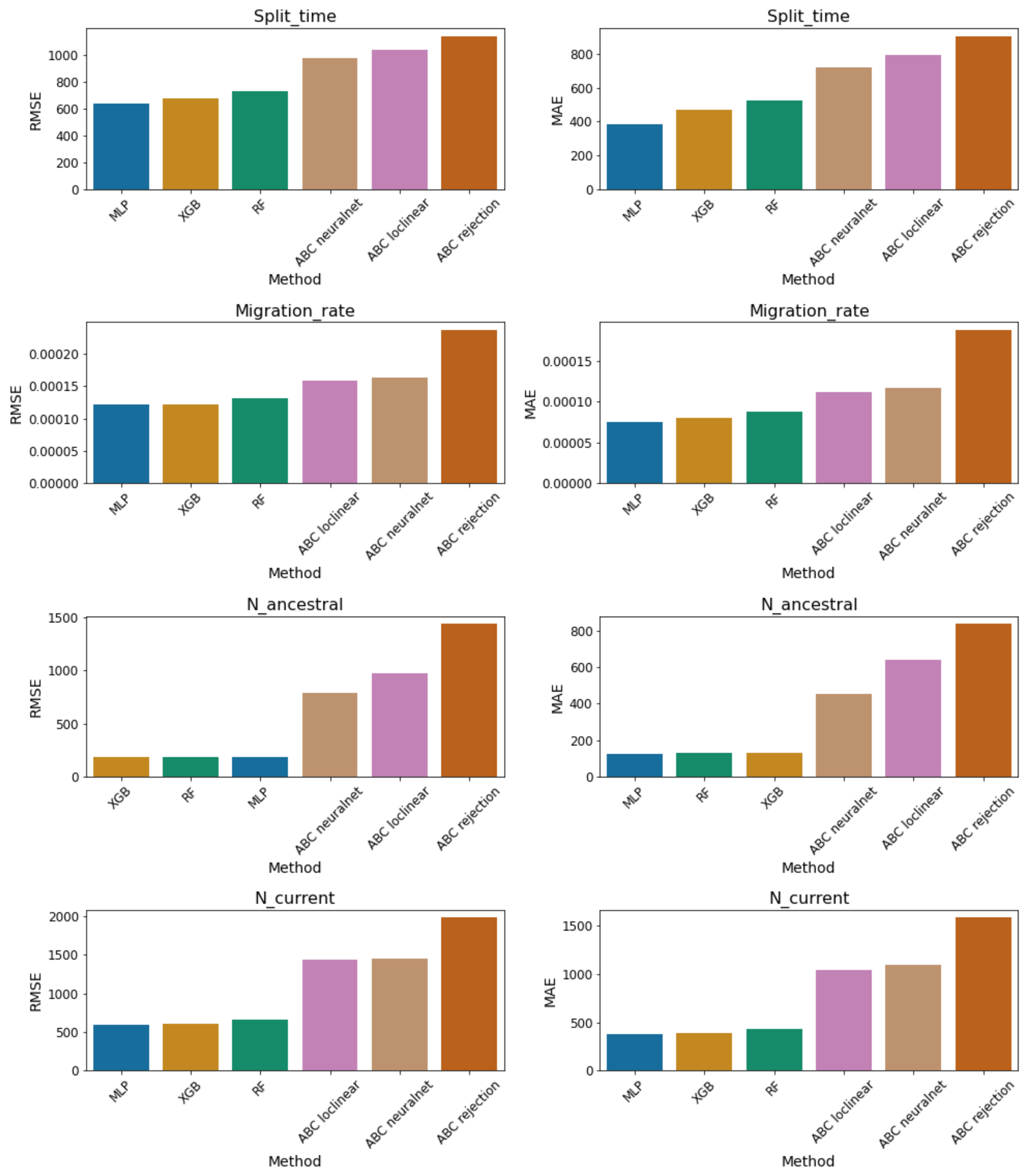


Figure S4. Comparisons of the Root Mean Square Error (RMSE) and Mean Absolute Error (MAE) of the three ML methods with the ABC algorithms. For each of the ABC algorithms we show the best performance in terms of tolerance threshold and choice of mean or median posterior as estimator.

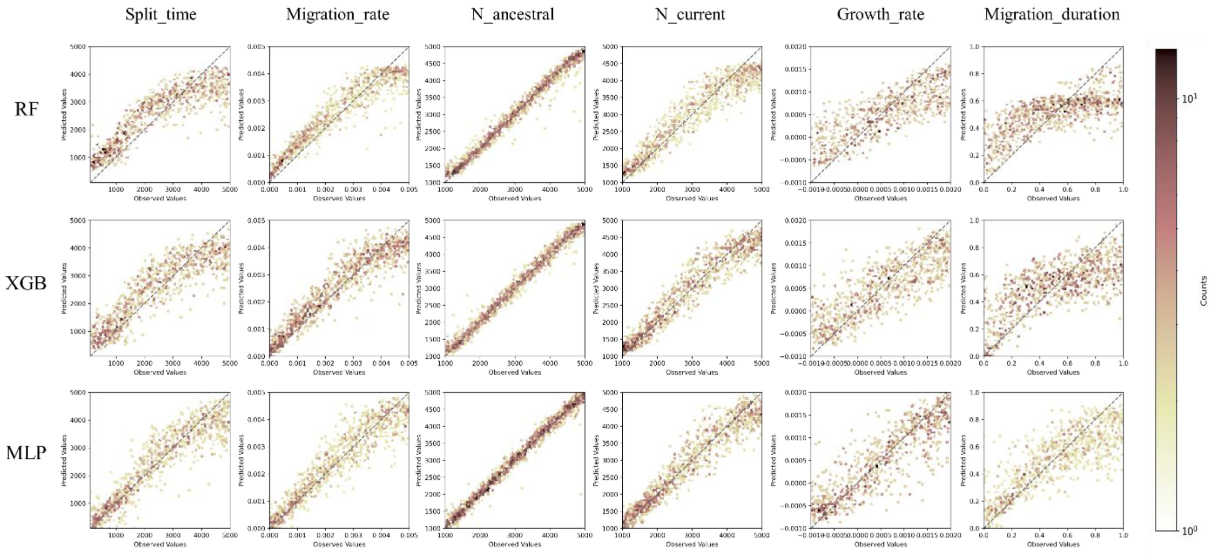


Figure S5. Predicted values versus observed values for the six secondary contact with varying population sizes parameters estimated with three different ML methods (RF: Random Forest, XGB: XGBoost, MLP: Multilayer perceptron).

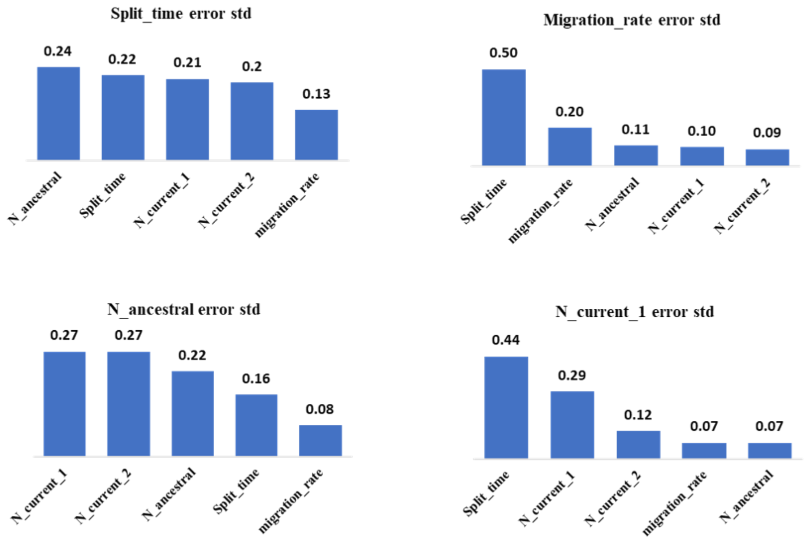


Figure S6. Feature importance for the prediction of the standardized errors of each demographic parameters modelled with a Random Forest based on all the demographic parameters.

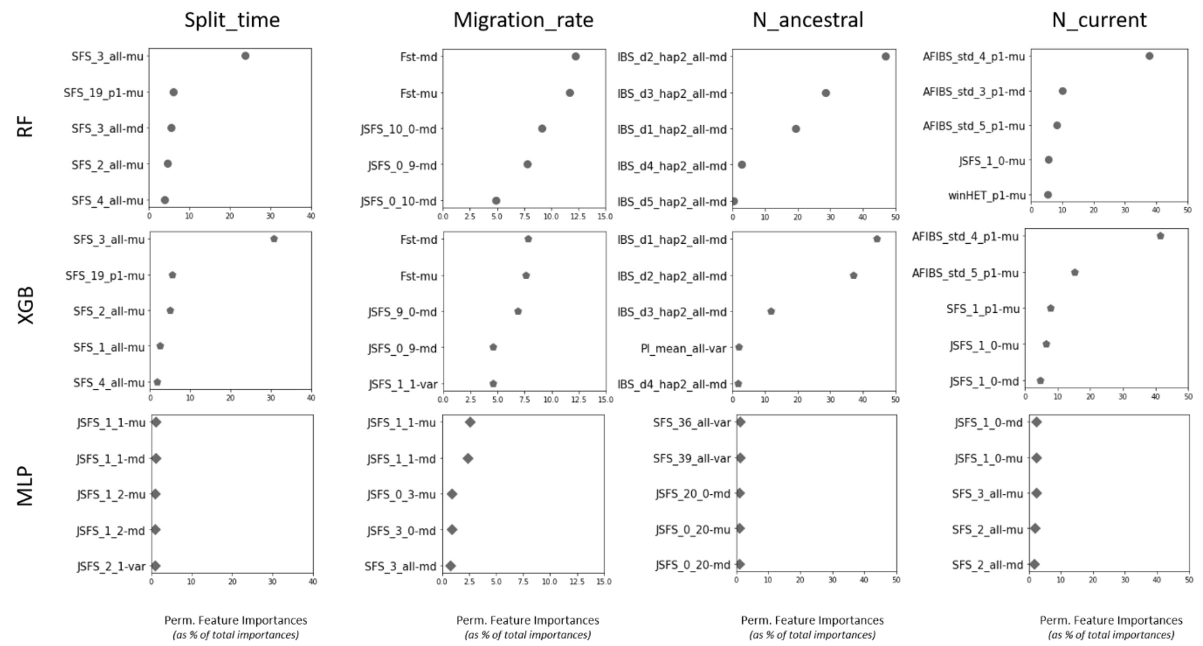


Figure S7. Top five summary statistics with the highest importance level for the three methods, computed with Permutation Feature Importance (PFI). The X-axis represents the percentage of the total PFI for the corresponding statistic.

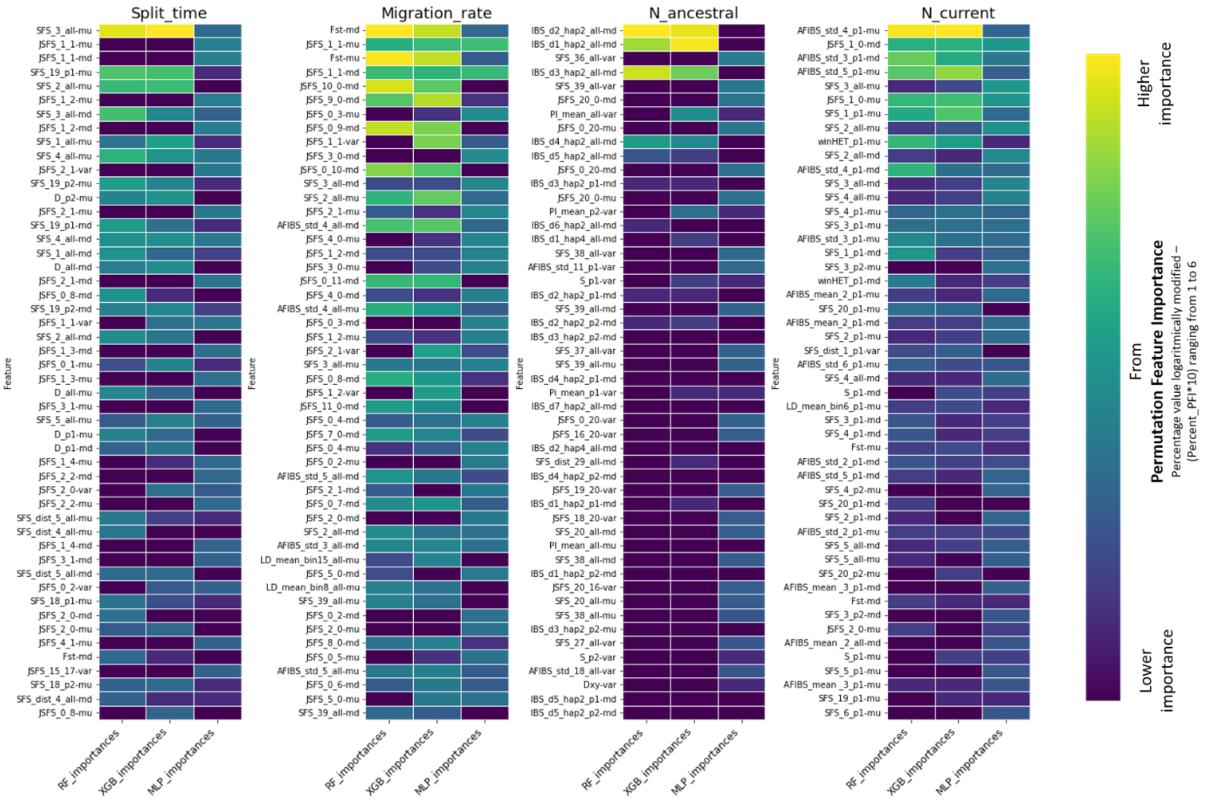


Figure S8. Importance levels of individual summary statistics for predicting demographic parameters using the three methods. We only displayed the first 50 variables in terms of highest rank of importance for one of the three methods.

| Class | Summary statistic | Description |
|---|---|---|
| AFIBS | AFIBS_***type_i_pop*** | ***type*** (mean or std) of the length of identical region around the focal SNP for frequency ***i*** of the derived allele for ***pop*** |
| D | D_***pop*** | Tajima's D computed for ***pop*** |
| Dxy | Dxy | genetic divergence |
| Fst | Fst | fixation index |
| IBS | IBS_***decile_hap_pop*** | decile ***decile*** of the length distribution of IBS segments shared between ***hap*** haplotypes for ***pop*** |
| JSFS | JSFS_***i_j*** | percentage of SNP where the derived allele is at frequency ***i*** in population 1 and ***j*** in population 2 |
| LD | LD_***type_bin_pop*** | ***type*** (mean or std) of $r^2$ for the length interval ***bin*** for ***pop*** |
| PI | PI_***type_pop*** | ***type*** (mean or std) of nucleotide diversity for ***pop*** |
| S | S_***pop*** | proportion of segregating sites for ***pop*** |
| SFS | SFS_(dist_)***i_pop*** | SFS_: Percentage of SNP where the derived allele is at frequency ***i*** in ***pop*** or SFS_ dist_: std of the distances separating two adjacent SNPs at frequency ***i*** for ***pop*** |
| winHET | winHet_***type_pop*** | ***type*** (mean or std) haplotypic diversity for ***pop*** |

**Table S1.** Classes of summary statistics. **pop**: "p1" for population 1, "p2" for population 2 and "all" for population 1 and 2. **-mu**, **-md** and **-var** at the end of the summary statistic is respectively for mean, median and variance across loci.

| Demographic parameters | RF | XGB | MLP |
|---|---|---|---|
| IM - Split_time | {300, 20, 10, 5} | {200, 0.2, 7} | {[2500, 1000, 500, 200, 80, 30, 10], relu} |
| IM - Migration_rate | {200, 15, 15, 10} | {200, 0.2, 6} | {[1000, 500, 200, 80, 30, 10], relu} |
| IM - N_ancestral | {200, 15, 15, 5} | {200, 0.1, 6} | {[2000, 1000, 500, 200, 80, 30, 10], relu} |
| IM - N_current | {200, 20, 15, 5} | {150, 0.1, 6} | {[2000, 1000, 500, 200, 80, 30, 10], relu} |
| SC - Split_time | {300, 20, 10, 5} | {200, 0.2, 7} | {[2500, 1000, 500, 200, 80, 30, 10], relu} |
| SC - Migration_rate | {200, 15, 15, 10} | {200, 0.2, 6} | {[1000, 500, 200, 80, 30, 10], relu} |
| SC - N_ancestral | {200, 15, 15, 5} | {200, 0.1, 6} | {[2000, 1000, 500, 200, 80, 30, 10], relu} |
| SC - N_current | {200, 20, 15, 5} | {150, 0.1, 6} | {[2000, 1000, 500, 200, 80, 30, 10], relu} |
| SC- Growth_rate | {200, 15, 15, 5} | {150, 0.2, 8} | {[2000, 1000, 500, 200, 80, 30], relu} |
| SC - Migration_duration | {200, 15, 15, 10} | {200, 0.2, 6} | {[2000, 1000, 500, 200, 80, 30, 10], relu} |

**Table S2.** Hyperparameters for the standard isolation with migration model (IM) and the secondary contact with varying population sizes model (SC). {n_estimators, max_depth, min_samples_split, min_samples_leaf} are indicated in the table for RF, {num_rounds, eta, max_depth} are indicated in the table for XGB, {Nb of neurons in hidden layers, activation function} are indicated in the table for MLP.

| Parameter | Method | Tolerance | ABC rejection | | ABC loclinear | | ABC neuralnet | |
|---|---|---|---|---|---|---|---|---|
| | | | MAE | RMSE | MAE | RMSE | MAE | RMSE |
| Split_time | median | 5.00E-04 | 9.76E+02 | 1.30E+03 | 1.46E+03 | 1.78E+03 | 7.60E+02 | 1.07E+03 |
| Split_time | mean | 5.00E-04 | 9.44E+02 | 1.23E+03 | 1.38E+03 | 1.68E+03 | 7.43E+02 | 1.02E+03 |
| Split_time | median | 0.001 | 9.24E+02 | 1.22E+03 | 1.11E+03 | 1.38E+03 | 7.33E+02 | 1.03E+03 |
| Split_time | mean | 0.001 | **9.06E+02** | 1.16E+03 | 1.07E+03 | 1.32E+03 | **7.22E+02** | **9.77E+02** |
| Split_time | median | 0.005 | 9.09E+02 | 1.16E+03 | 8.92E+02 | 1.12E+03 | 7.37E+02 | 1.00E+03 |
| Split_time | mean | 0.005 | 9.17E+02 | **1.14E+03** | 9.15E+02 | 1.12E+03 | 7.54E+02 | 9.92E+02 |
| Split_time | median | 0.01 | 9.38E+02 | 1.17E+03 | **7.94E+02** | **1.04E+03** | 7.61E+02 | 1.02E+03 |
| Split_time | mean | 0.01 | 9.52E+02 | 1.16E+03 | 8.17E+02 | 1.04E+03 | 7.84E+02 | 1.02E+03 |
| Split_time | median | 0.02 | 9.79E+02 | 1.21E+03 | 8.14E+02 | 1.06E+03 | 8.04E+02 | 1.06E+03 |
| Split_time | mean | 0.02 | 9.97E+02 | 1.20E+03 | 8.45E+02 | 1.06E+03 | 8.36E+02 | 1.05E+03 |
| Split_time | median | 0.05 | 1.06E+03 | 1.27E+03 | 8.93E+02 | 1.12E+03 | 8.93E+02 | 1.12E+03 |
| Split_time | mean | 0.05 | 1.08E+03 | 1.27E+03 | 9.34E+02 | 1.13E+03 | 9.33E+02 | 1.13E+03 |
| Migration_rate | median | 5.00E-04 | 1.99E-04 | 2.61E-04 | 2.95E-04 | 3.59E-04 | 1.22E-04 | 1.72E-04 |
| Migration_rate | mean | 5.00E-04 | 1.94E-04 | 2.49E-04 | 2.84E-04 | 3.44E-04 | 1.20E-04 | 1.68E-04 |
| Migration_rate | median | 0.001 | 1.92E-04 | 2.49E-04 | 1.92E-04 | 2.46E-04 | 1.18E-04 | 1.67E-04 |
| Migration_rate | mean | 0.001 | **1.88E-04** | 2.39E-04 | 1.89E-04 | 2.38E-04 | **1.17E-04** | **1.63E-04** |
| Migration_rate | median | 0.005 | 1.92E-04 | 2.42E-04 | **1.12E-04** | 1.59E-04 | 1.24E-04 | 1.72E-04 |
| Migration_rate | mean | 0.005 | 1.92E-04 | **2.38E-04** | 1.15E-04 | **1.59E-04** | 1.27E-04 | 1.70E-04 |
| Migration_rate | median | 0.01 | 1.94E-04 | 2.43E-04 | 1.23E-04 | 1.68E-04 | 1.31E-04 | 1.79E-04 |
| Migration_rate | mean | 0.01 | 1.95E-04 | 2.40E-04 | 1.27E-04 | 1.70E-04 | 1.34E-04 | 1.77E-04 |
| Migration_rate | median | 0.02 | 1.98E-04 | 2.46E-04 | 1.39E-04 | 1.87E-04 | 1.41E-04 | 1.89E-04 |
| Migration_rate | mean | 0.02 | 1.99E-04 | 2.44E-04 | 1.44E-04 | 1.87E-04 | 1.45E-04 | 1.88E-04 |
| Migration_rate | median | 0.05 | 2.05E-04 | 2.51E-04 | 1.59E-04 | 2.08E-04 | 1.59E-04 | 2.08E-04 |
| Migration_rate | mean | 0.05 | 2.08E-04 | 2.50E-04 | 1.63E-04 | 2.06E-04 | 1.63E-04 | 2.06E-04 |
| N_ancestral | median | 5.00E-04 | 9.06E+02 | 1.57E+03 | 2.93E+03 | 3.60E+03 | 4.92E+02 | 8.36E+02 |
| N_ancestral | mean | 5.00E-04 | 8.69E+02 | 1.49E+03 | 2.82E+03 | 3.43E+03 | 4.68E+02 | 7.91E+02 |
| N_ancestral | median | 0.001 | 8.70E+02 | 1.52E+03 | 1.85E+03 | 2.38E+03 | 4.85E+02 | 8.34E+02 |
| N_ancestral | mean | 0.001 | **8.39E+02** | **1.44E+03** | 1.83E+03 | 2.28E+03 | **4.56E+02** | **7.86E+02** |
| N_ancestral | median | 0.005 | 8.70E+02 | 1.51E+03 | 9.29E+02 | 1.23E+03 | 4.96E+02 | 8.47E+02 |
| N_ancestral | mean | 0.005 | 8.64E+02 | 1.44E+03 | 9.59E+02 | 1.23E+03 | 4.86E+02 | 8.22E+02 |
| N_ancestral | median | 0.01 | 9.20E+02 | 1.56E+03 | 6.74E+02 | **9.67E+02** | 5.24E+02 | 8.73E+02 |
| N_ancestral | mean | 0.01 | 9.21E+02 | 1.49E+03 | 6.96E+02 | 9.80E+02 | 5.22E+02 | 8.62E+02 |
| N_ancestral | median | 0.02 | 1.05E+03 | 1.72E+03 | **6.43E+02** | 9.92E+02 | 5.65E+02 | 9.23E+02 |
| N_ancestral | mean | 0.02 | 1.03E+03 | 1.61E+03 | 6.59E+02 | 1.00E+03 | 5.75E+02 | 9.27E+02 |
| N_ancestral | median | 0.05 | 1.25E+03 | 1.98E+03 | 6.64E+02 | 1.03E+03 | 6.62E+02 | 1.03E+03 |
| N_ancestral | mean | 0.05 | 1.22E+03 | 1.83E+03 | 6.82E+02 | 1.06E+03 | 6.80E+02 | 1.05E+03 |
| N_current | median | 5.00E-04 | 1.67E+03 | 2.20E+03 | 2.80E+03 | 3.39E+03 | 1.14E+03 | 1.54E+03 |
| N_current | mean | 5.00E-04 | 1.65E+03 | 2.12E+03 | 2.68E+03 | 3.22E+03 | 1.14E+03 | 1.51E+03 |
| N_current | median | 0.001 | 1.59E+03 | 2.07E+03 | 1.54E+03 | 2.13E+03 | **1.09E+03** | 1.46E+03 |
| N_current | mean | 0.001 | 1.62E+03 | 2.02E+03 | 1.51E+03 | 2.02E+03 | 1.12E+03 | **1.45E+03** |
| N_current | median | 0.005 | **1.58E+03** | **1.99E+03** | **1.04E+03** | **1.44E+03** | 1.16E+03 | 1.49E+03 |
| N_current | mean | 0.005 | 1.69E+03 | 2.03E+03 | 1.09E+03 | 1.46E+03 | 1.27E+03 | 1.56E+03 |
| N_current | median | 0.01 | 1.64E+03 | 2.02E+03 | 1.27E+03 | 1.64E+03 | 1.24E+03 | 1.57E+03 |
| N_current | mean | 0.01 | 1.75E+03 | 2.08E+03 | 1.38E+03 | 1.71E+03 | 1.37E+03 | 1.66E+03 |
| N_current | median | 0.02 | 1.72E+03 | 2.08E+03 | 1.29E+03 | 1.61E+03 | 1.36E+03 | 1.68E+03 |
| N_current | mean | 0.02 | 1.84E+03 | 2.15E+03 | 1.42E+03 | 1.71E+03 | 1.51E+03 | 1.79E+03 |
| N_current | median | 0.05 | 1.85E+03 | 2.19E+03 | 1.57E+03 | 1.90E+03 | 1.57E+03 | 1.89E+03 |
| N_current | mean | 0.05 | 1.96E+03 | 2.27E+03 | 1.73E+03 | 2.01E+03 | 1.73E+03 | 2.01E+03 |

**Table S3.** Mean Absolute Errors (MAE) and Root Mean Square Errors (RMSE) for the demographic parameters obtained for the three ABC algorithms, based on the mean posterior and the median posterior, and for tolerance levels ranging from $5\times10^{-4}$ to $5\times10^{-2}$.

| Method | RMSE | MAE *(se)* | NMAE *(se)* |
| --- | --- | --- | --- |
| RF | 6.43E+02 | 4.13E+02 *(8.26)* | 4.18E-02 *(8.35E-04)* |
| XGB | 5.90E+02 | 3.78E+02 *(7.57)* | 3.82E-02 *(7.65E-04)* |
| MLP | **5.75E+02** | **3.68E+02** *(7.35)* | **3.71E-02** *(7.43E-04)* |

**Table S4.** Test scores of the three ML methods for predicting the N_current_2 parameter.
